# Chemical Impacts of the Microbiome Across Scales Reveal Novel Conjugated Bile Acids

**DOI:** 10.1101/654756

**Authors:** Robert A. Quinn, Alison Vrbanac, Alexey V. Melnik, Kathryn A. Patras, Mitchell Christy, Andrew T. Nelson, Alexander Aksenov, Anupriya Tripathi, Greg Humphrey, Ricardo da Silva, Robert Bussell, Taren Thron, Mingxun Wang, Fernando Vargas, Julia M. Gauglitz, Michael J. Meehan, Orit Poulsen, Brigid S. Boland, John T. Chang, William J. Sandborn, Meerana Lim, Neha Garg, Julie Lumeng, Barbara I. Kazmierczak, Ruchi Jain, Marie Egan, Kyung E. Rhee, Gabriel G. Haddad, Dionicio Siegel, Sarkis Mazmanian, Victor Nizet, Rob Knight, Pieter C. Dorrestein

**Affiliations:** Skaggs School of Pharmacy and Pharmaceutical Sciences, University of California San Diego, La Jolla, CA; Center for Microbiome Innovation, University of California San Diego, La Jolla, CA; Department of Pediatrics, University of California San Diego, La Jolla, CA; Department of Radiology, University of California San Diego, La Jolla, CA; Division of Biology and Biological Engineering, California Institute of Technology, Pasadena, CA; Division of Gastroenterology, Department of Medicine, University of California San Diego, La Jolla, CA; School of Chemistry and Biochemistry, Georgia Institute of Technology, Atlanta, GA; Emory-Children’s Cystic Fibrosis Center, Atlanta, GA; Department of Pediatrics, University of Michigan, Ann Arbor, MI; Department of Internal Medicine, Yale School of Medicine, New Haven, CT; Department of Pediatrics, Yale School of Medicine, New Haven, CT; Department of Computer Science and Engineering, University of California San Diego, La Jolla, CA

## Abstract

A mosaic of cross-phyla chemical interactions occurs between all metazoans and their microbiomes. In humans, the gut harbors the heaviest microbial load, but many organs, particularly those with a mucosal surface, associate with highly adapted and evolved microbial consortia^1^. The microbial residents within these organ systems are increasingly well characterized, yielding a good understanding of human microbiome composition, but we have yet to elucidate the full chemical impact the microbiome exerts on an animal and the breadth of the chemical diversity it contributes^2^. A number of molecular families are known to be shaped by the microbiome including short-chain fatty acids, indoles, aromatic amino acid metabolites, complex polysaccharides, and host lipids; such as sphingolipids and bile acids^3–11^. These metabolites profoundly affect host physiology and are being explored for their roles in both health and disease. Considering the diversity of the human microbiome, numbering over 40,000 operational taxonomic units^12^, a plethora of molecular diversity remains to be discovered. Here, we use unique mass spectrometry informatics approaches and data mapping onto a murine 3D-model^13–15^ to provide an untargeted assessment of the chemical diversity between germ-free (GF) and colonized mice (specific-pathogen free, SPF), and report the finding of novel bile acids produced by the microbiome in both mice and humans that have evaded characterization despite 170 years of research on bile acid chemistry^16^.

## Main

In total, 96 sample sites, covering 29 organs, producing 768 samples (excluding controls, Fig. S1) were analyzed from four GF and four colonized mice by LC-MS/MS mass spectrometry and 16S rRNA gene sequencing. The metabolome data was most strongly influenced by organ source, but as expected, the microbiome was dictated by colonization status (Fig. 1a,b). GF mice and sterile organs in SPF mice clustered tightly with background sequence reads from blanks (reflecting their sterility), whereas colonized organs within the SPF mice clustered apart from these samples (Fig. 1a,b). Mapping the principle coordinate values of the two data types onto the murine 3-D model showed how the gut samples were similar, but important differences were observed, including separation of the stool sample from the upper GI tract in the metabolome but not in the microbiome, and similarity between the esophageal and gut microbiomes. The strongest separation in the metabolome between colonization states was present in the stool, cecum, other regions of the GI tract, and samples from the surface of the animals including ears and feet (Fig. 1c). The liver also had signatures suggestive of metabolomic differences between the GF and SPF mice, but these were not significant compared to the within individual variation (Fig. 1, Fig. S2).

**Figure 1.**
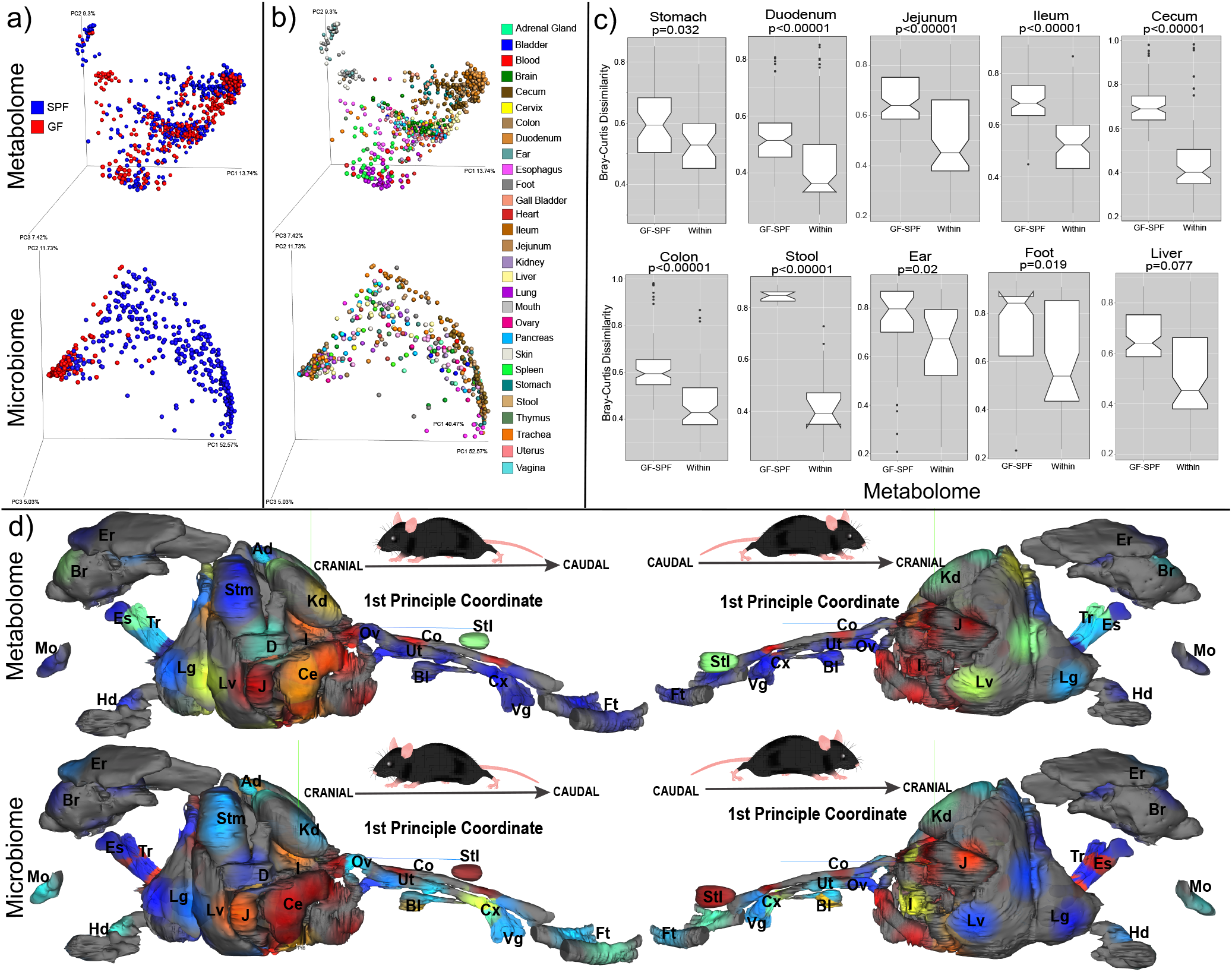
a) Principal coordinate analysis (PCoA) of microbiome and mass spectrometry data highlighted by sample source as GF or SPF. b) Same data highlighted by organ source. c) Bray-Curtis dissimilarities of the metabolome data collected from murine organs. The dissimilarities are calculated within individual mice of the same group (GF or SPF, “Within”) or across the GF and SPF groups (GF-SPF). Organs with multiple samples are pooled, but only samples collected from exact same location are compared. d) 3-D model of murine organs mapped with the mean 1^st^ principle coordinate value from the four GF and four SPF mice. High values across the 1^st^ PC are shown in red and lower values are shown in blue. The PC1 values are from the data in panels a) and b). (Er=ear, Br=brain, Ad=adrenal gland, Es=esophagus, Tr=trachea, Stm=stomach, Kd=kidney, Mo=mouth, D=duodenum, Ov=ovary, Co=colon, Stl=stool, Hd=hand, Lg=lung, Lv=liver, J=jejunum, Ce=cecum, Bl-bladder, Ut=uterus, Cx=cervix, Vg=vagina, Ft=feet)

Molecular networking is a novel spectral alignment algorithm that enables identification of unique molecules in mass spectrometry data and the relationships between related spectra^14^. Applying molecular networking to this comprehensive murine dataset identified 7,913 unique spectra (representing putative molecules) of which 14.7% were exclusively observed in colonized mice and 10.0% were exclusive to GF (Fig. 2). Although the overall profiles exhibited the strongest difference in the GI tract, molecular networking showed that all organs had some unique molecular signatures from the microbiome, ranging from 2% in the bladder to 44% in stool (Fig. 2). As expected, the metabolome of the cecum, site of microbial fermentation of food products, was profoundly affected by the microbiota, but other GI sites had weaker signatures. Spectral library searching enabled annotation of 8.86% of nodes in the molecular network (n=700 annotated nodes^13,17^); which included members of the molecular families of plant products, such as soyasaponins and isoflavonoids (sourced from the soybean (*Glycine max*, f. *Fabaceae*) component of mouse chow), host lipids and microbial metabolic products (Fig. 2a). Many of the unique signatures attributed to the microbiome were the result of metabolism of plant triterpenoids and flavonoids from food (Supplemental Data, Fig. S3, S4). These effects were location specific, indicating that the microbiome inhabits spatially distinct and varied niche space throughout an organism, exerting location-dependent effects on host physiology through the metabolism of xenobiotics and modification of host molecules.

**Figure 2.**
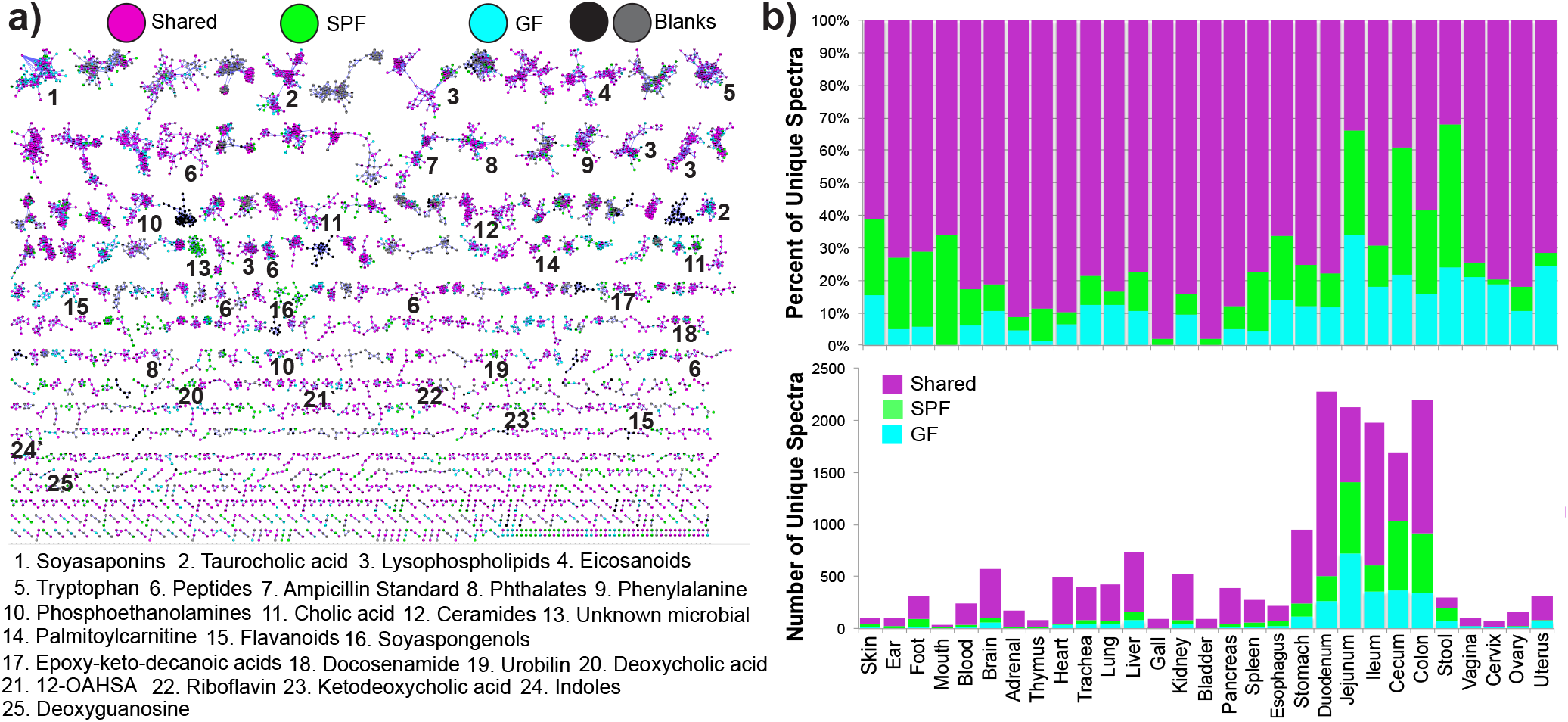
a) Molecular network of LC-MS/MS data with nodes colored by source as GF, SPF, shared, or detected in blanks. Molecular families with metabolites annotated by spectral matching in GNPS are listed by a number corresponding to the molecular family. These are level 2 or 3 annotations according to the metabolomics standards consortium ^31^. b) percentage of total nodes from each organ sourced from GF only, SPF only or shared and the total number of unique nodes from each murine class per organ.

The strong impacts from the microbiome in the gastrointestinal (GI) tract led to deeper analysis of the molecular changes in this organ system. A random forests classification was used to identify the most differentially abundant molecules between the GF and SPF GI tracts. The metabolome of both the GF and SPF mice changed through the different sections of the digestive system (Fig. 3a). While changes through the upper GI tract were subtle in GF mice, SPF animals had progressive transitions in this region (Fig. 3a). A major transition occurred between the ileum and cecum in both groups, but the specific molecules that were changing were different between them (Fig. 3a). Many unique metabolites in SPF mice were unknown compounds, but known molecules were also identified including bile acids and soyasaponins (Fig. 3a, Supplementary Data, Fig. S3,S5). The Shannon diversity index of the GF and SPF mouse metabolome was mirrored in the upper GI tract, both being low in the esophagus and higher in the stomach and duodenum, however, upon transition to the cecum, the diversity of the two groups of mice began to separate (Fig. 3c,d). The molecular diversity in the cecum and colon of colonized mice was significantly higher than GF mice (Mann-Whitney U-test), but not in the stool samples (Fig. 3c).

**Figure 3.**
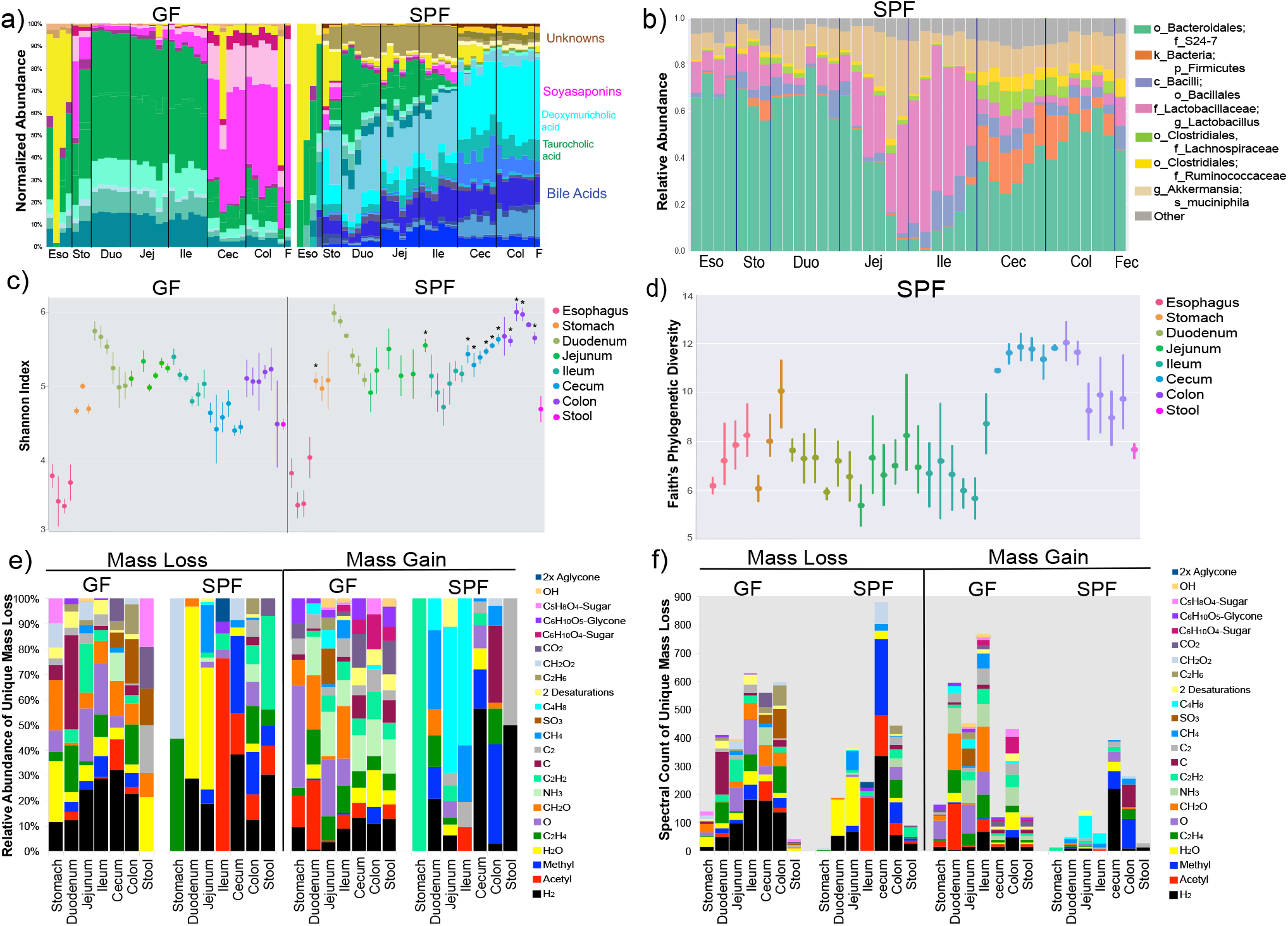
a) Mean normalized abundance of the top 30 most differentially abundant metabolites between GF and SPF mice. The metabolites are colored according to molecular family, where bile acids are green and blue, respectively, soyasaponins are pink and unknown molecules are brown/yellow. Colors corresponding to taurocholic acid (green) and deoxymuricholic acid (teal) are highlighted for reference. b) Microbiome of the murine GI tract in SPF mice. Taxa of relevance are color coded according to the legend. c) Mean and 95% confidence interval of the Shannon-Weiner diversity of the metabolomic data in each GI tract sample for GF and SPF mice. Statistical significance between metabolome diversity in the same sample location between GF and SPF mice was tested with the Mann-Whitney U-test (*=p<0.05). d) Mean Faith’s phylogenetic diversity (with 95% confidence interval) of the microbiome through the SPF GI tract. e) Results of meta-mass shift chemical profiling ^19^ showing the relative abundance of the parent mass differences between unique nodes in either GF or SPF mice to the total. Each mass difference corresponds to the node-to-node gain or loss of a particular chemical group. f) Counts of the number of mass shifts of the parent mass differences between nodes showing where the most abundant molecular

We also compared the changing microbial community through the GI tract in the context of the changes observed in the molecular data. Similar to the metabolites, microbiome transitions were observed traversing the GI tract (Fig. 3b). The corresponding microbial diversity of the colonized animals showed a similar profile to the metabolome, mostly stable through the upper parts of the system and then abruptly increasing at the cecum, followed by a decrease in the colon and stool (Fig. 3d). However, an interesting contrast was observed where a high diversity of the metabolome in the duodenum corresponded to a lower microbial diversity. We hypothesize that this contrasting result was due to the secretion of bile acids from the gallbladder at this location. Because these molecules possess antimicrobial properties, their high abundance may explain the lower microbial diversity in the upper GI tract^18^, while simultaneously, microbial modification of the molecules increases their molecular diversity. After the duodenum, changes in the diversity of microbiome and metabolome were closely aligned, but colonized mice had greater molecular diversity in the cecum and colon. This shows that microbial activity in these organs was altering the molecules present, particularly bile acids, soyasaponins, flavonoids, and other unknown compounds, which expanded the metabolomic diversity of the cecum (supplementary results).

Molecular networking also enabled meta-mass shift chemical profiling^19^ of the GF and SPF GI tract, which is an analysis of chemical transformations based on parent mass shifts between related nodes without the requirement of knowing the molecular structure. For example, a unique node found in colonized mice with an 18.015 Da difference represents H_2_O and 2.016 Da is H_2_. In colonized animals, there was a strong signature for the loss of water in the duodenum and jejunum and the loss of H_2_, acetyl and methyl groups in latter parts of the GI tract (Fig. 3e,f). GF mice had notable mass gains corresponding to monosaccharides in all regions of the GI tract, which were absent in SPF animals. Instead, a mass gain of C_4_H_8_ was seen in the jejunum and ileum of SPF mice, which was associated with the conjugated bile acid glycocholic acid (Fig. 3e,f). A significant portion of the dehydrogenation and dehydroxylation mass shifts from the microbiome were associated with bile acids, indicating that microbial enzymes acted on C-C double bonds of the cholic acid backbone and removed hydroxyl groups, which is a known microbial transformation^3^. Deacetylations were also prevalent in SPF animals, though the metabolites upon which these losses were occurring remain mostly unidentified. Overall, both GF and SPF mice had many cases of mass loss between related molecules, but there were comparably fewer molecules in the colonized mice that showed gain of a molecular group (Fig. 3f). This indicates that the microbiome contributes more to the catabolic breakdown of molecules and less to anabolism; however, one interesting anabolic reaction that was detected was the addition of C_4_H_8_ on glycocholic acid, which we subsequently investigated further.

Glycine and taurine conjugated bile acids were detected in both GF and SPF mice. As they moved through the GI tract, the conjugated amino acid was removed in SPF mice only, representing a known microbial transformation (Fig. S5,^20^). In the bile acid molecular network that contained taurocholic acid and glycocholic acid there were modified forms of these compounds that were only present in colonized animals. These nodes were related to the glycocholic acid through spectral similarity and to the sulfated form (Fig. 4a) and one of them corresponded to the addition of C_4_H_8_ described above. Analysis of the MS/MS spectra of the three nodes *m*/*z* 556.363, *m*/*z* 572.358 and *m*/*z* 522.379 showed maintenance of the core cholic acid, but with a fragmentation pattern characteristic of the presence of the amino acids phenylalanine, tyrosine and leucine through an amide bond at the conjugation site in place of glycine or taurine (Fig. S6). In the extensive bile acid literature, representing 170 years of bile acid structural analysis and greater than 42,000 publication records in PubMed, the only known conjugations of murine (and human) bile acids were those of glycine and taurine^16^. Here, we have found a set of unique amino acid conjugations to cholic acid mediated by the microbiome creating the novel bile acids phenylalanocholic acid, tyrosocholic acid and leucocholic acid. These structures were validated with synthesized standards using NMR and mass spectrometry methods (Supplemental methods and Fig. S7, S8, S9, S10, S11). These uniquely conjugated bile acids were detected in the duodenum, jejunum and ileum of SPF mice, with phenylalanocholic acid being the most abundant (Fig. 4). In comparison, glycocholic acid was present in the latter parts of the GI tract (cecum and colon), whereas taurocholic acid was most abundant in the upper parts of the GI tract (reduced through the lower GI tract in SPF mice). The concentration of phenylalanocholic acid in mouse ileal content from the four mice was 0.59 μM (s.d. 0.21) in the duodenum, 3.0 μM (s.d. 4.43) in the jejunum and 5.25 μM (s.d. 2.42) in the ileum, with its highest concentration reaching 13.24 μM in a single jejunum sample (Fig. S12). These findings demonstrate that these novel amino acid conjugates are abundant in the upper GI tract of mice on a normal soy-based diet and require the microbiota for their production, but were subsequently absorbed, further modified, or deconjugated again upon travel to the cecum.

**Figure 4.**
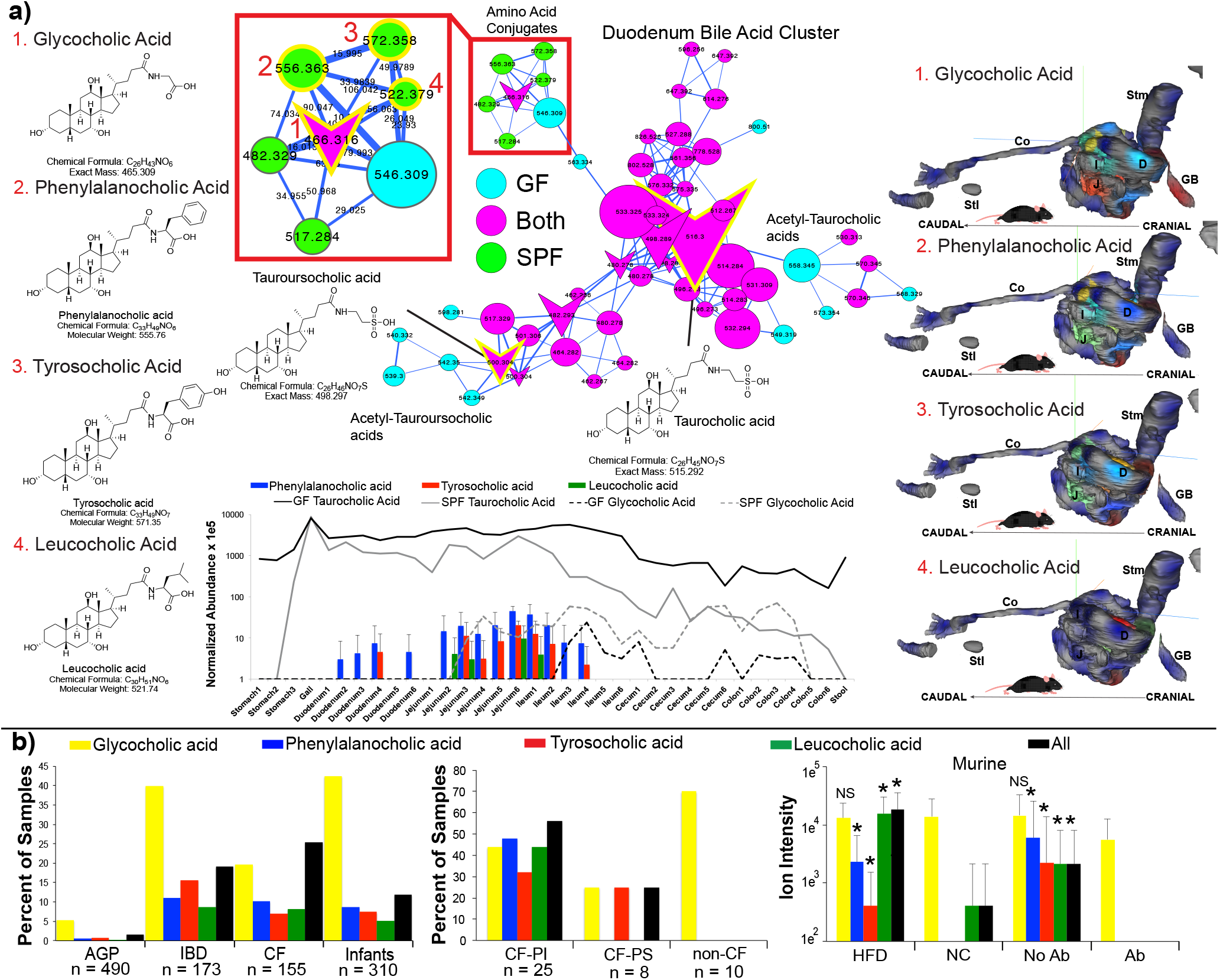
a) Structures, molecular network, 3D-molecular cartography and abundance through GI tract of novel microbiome associated bile acids in this murine study. Structures of the previously known conjugates glycocholic acid, taroursocholic acid and taurocholic acid are shown for comparison to structures of the newly discovered amino acid conjugates. The molecular network of these bile acids is shown with mapping to the GF and SPF mice according to the color legend. An inset highlighting the parent masses and mass differences between the newly discovered molecules is shown for clarity. 3D-molecular cartography maps the mean abundance and standard deviations of the mean of the newly discovered conjugates onto a 3D-rendered model of the murine GI tract and the relative abundances of the molecules through the GI tract samples compared to the host produced glycocholic acid and taurocholic acid are also shown. b) Bar plots of the percent of samples positive for the novel bile acids from publically available datasets on GNPS. Percent of patients where novel bile acids were detected from two human studies of cystic fibrosis patients compared to non-CF controls. Comparison of the abundance of novel conjugates in a controlled murine study previously published where animals fed high fat diet (HFD) or normal chow (NC) were compared and those treated with antibiotics ^22^. AGP = American Gut Project ^21^, IBD = Inflammatory Bowel Disease, CF = cystic fibrosis, PI = pancreatic insufficient, PS = pancreatic sufficient.

Because GNPS is a public repository of mass spectrometry data from a wide variety of biological systems, we used an analysis feature called “single spectrum search” to search all 739 publically available data sets for the presence of MS/MS spectra matching these conjugated bile acids (April 27, 2018,^13^). Spectral matches corresponding to phenylalanocholic acid, tyrosocholic acid and leucocholic acid were found in 19 other studies comprising samples from the GI tract of both mice (with at least one conjugate found in 3.2 to 59.4% of all samples, Fig. S13) and humans (in 1.6 to 25.3% of all samples, Fig. S13). In a crowd-sourced fecal microbiome and metabolome study at least one of these unique bile acids was found in 1.6% of human fecal samples with tyrosocholic acid being the most prevalent (n=490, the American Gut Project ^21^, Fig. 4b). They were found in higher frequency in fecal samples collected without swabs, including studies of patients with inflammatory bowel syndrome, cystic fibrosis (CF) and infants (Fig. 4b). Re-analysis of data from a previously published study of the murine microbiome and liver cancer enabled a comparison of the abundance of these molecules in mice fed a high-fat-diet (HFD) and treated with antibiotics^22^, Fig. 4b). Supporting the role of the microbiome in their production, the Phe/Tyr/Leu amino acid conjugates were decreased with antibiotic exposure, whereas glycocholic acid, which is synthesized by host liver enzymes, was not. In contrast, these microbial bile acids were more abundant in mice fed HFD, with no change observed in the host conjugated glycocholic acid^22^. In a separate data set where atherosclerosis-prone mice were similarly fed a HFD the novel conjugates were also increased over time, but not on normal chow and the host-conjugated taurocholic acid did not change significantly (Fig. S14). Finally, exploration into the metadata associated with a public study of a pediatric CF patient cohort showed that there was a higher prevalence of these compounds in CF patients compared to healthy controls, particularly those with pancreatic insufficiency (Fig. 4b). Insufficient production of pancreatic lipase in the CF gut results in the buildup of fat and a microbial dysbiosis^23^, which parallels the gut microbial ecosystem in mice fed HFD.

The first chemical characterization of a bile acid was in 1848^24^, the first correct structure of a bile acid related molecule was elucidated in 1932^25^ and bile acid metabolism by the microbiome has been known since the 1960s^26^. Since then, microbial alteration of bile acids has been known to occur through four principal mechanisms: dehydroxylation, dehydration and epimerization of the cholesterol backbone, and deconjugation of the amino acids taurine or glycine^3,27,28^. Here, using a simple experiment with colonized and sterile mice, we have identified a fifth mechanism of bile acid transformation by the microbiome mediated by a completely novel mechanism: conjugation of the cholesterol backbone with the amino acids phenylalanine, leucine and tyrosine. Further research is required to determine the microbial producers of these compounds and their role in gut microbial ecology, especially considering the important findings that microbiome based bile acid metabolism can affect *C. difficile* infections^29^ or regulate liver cancer^30^. The findings reported here show that all bile acid research to date have overlooked a significant component of the human bile acid pool produced by the microbiome.

In conclusion, the chemistry of all major organs and organ systems are affected by the presence of a microbiome. The strongest signatures come from the gut through the modification of host bile acids and xenobiotics, particularly the breakdown of plant natural products from food. Addition of chemical groups to host molecules were more rare, but those that were detected were sourced from a unique alteration of host bile acids by the microbiome that changes our understanding of human bile after 170 years of research^16^. As the connections between us and our microbial symbionts becomes more and more obvious, a combination of globally untargeted approaches and the development of tools that interlink these data sets will enable us to identify novel molecules, leading to a better understanding of the deep connection between our microbiota and our health.

## Acknowledgements

The authors would like to acknowledge funding from the National Institutes of Health under project 5U01AI124316-03, 1R03CA211211-01, 1R01HL116235 and R01HD084163. Additionally, B.S.B. was supported by UCSD KL2 (1KL2TR001444), J.L. by grant 13EIA14660045 from the American Heart Association. We would like to acknowledge Gail Ackerman for her contributions to curate the metadata. We further thank Alan Hofmann and Lee Hagey for their insightful discussions on structural characterization of bile acids.

## Data Availability

All metabolomics data is available at GNPS (gnps.ucsd.edu) under the MassIVE id numbers: MSV000079949 (GF and SPF mouse data). Additional sample data: MSV000082480, MSV000082467, MSV000079134, MSV000082406. The sequencing data for the GF and SPF mouse study is available on the Qiita microbiome data analysis platform at Qiita.ucsd.edu under study ID 10801 and through the European Bioinformatics Institute accession number ERP109688.

## Contributions

PCD, RK and RQ designed the project

RQ, AA, AM, FV, JG, NG, AT, MC, ATN, MM, GH, RdS, and RB generated data

RQ, AV, AT and MC analyzed data

RQ, BB, ML, OP, JC, ML, JL, KP, BK, RJ, ME, KR, GH, KR, GC, WS and RB collected samples

PCD, RK, SM, VN and DS guided experimental design and analysis.

MW converted the data in GNPS, developed spectral search and molecular explorer.

TT, VN and SM raised animals and guided experimental design.

RQ and PD wrote the manuscript

## References

1. Consortium, H. Structure, function and diversity of the healthy human microbiome. Nature 486, 207–14 (2012).

2. Dorrestein, P. C., Mazmanian, S. K. & Knight, R. Finding the Missing Links among Metabolites, Microbes, and the Host. Immunity 40, 824–832 (2014).

3. Ridlon, J. M., Kang, D. J., Hylemon, P. B. & Bajaj, J. S. Bile acids and the gut microbiome. Curr. Opin. Gastroenterol. 30, 332–8 (2014).

4. Gilbert, J. A. et al. Microbiome-wide association studies link dynamic microbial consortia to disease. Nature 535, (2016).

5. Wikoff, W. R. et al. Metabolomics analysis reveals large effects of gut microflora on mammalian blood metabolites. Proc. Natl. Acad. Sci. U. S. A. 106, 3698–703 (2009).

6. Marcobal, A. et al. Metabolome progression during early gut microbial colonization of gnotobiotic mice. Sci. Rep. 5, 11589 (2015).

7. Miller, T. L. & Wolin, M. J. Pathways of acetate, propionate, and butyrate formation by the human fecal microbial flora. Appl. Environ. Microbiol. 62, 1589–92 (1996).

8. Gillner, M., Bergman, J., Cambillau, C., Fernström, B. & Gustafsson, J. A. Interactions of indoles with specific binding sites for 2,3,7,8-tetrachlorodibenzo-p-dioxin in rat liver. Mol. Pharmacol. 28, 357–63 (1985).

9. Martin, F.-P. J. et al. A top-down systems biology view of microbiome-mammalian metabolic interactions in a mouse model. Mol. Syst. Biol. 3, 112 (2007).

10. Moriya, T., Satomi, Y., Murata, S., Sawada, H. & Kobayashi, H. Effect of gut microbiota on host whole metabolome. Metabolomics 13, 101 (2017).

11. Swann, J. R. et al. Systemic gut microbial modulation of bile acid metabolism in host tissue compartments. Proc. Natl. Acad. Sci. U. S. A. 108 Suppl 1, 4523–30 (2011).

12. Structure, function and diversity of the healthy human microbiome. Nature 486, 207–14 (2012).

13. Wang, M. et al. Sharing and community curation of mass spectrometry data with Global Natural Products Social Molecular Networking. Nat. Biotechnol. 34, 828–837 (2016).

14. Watrous, J. et al. Mass spectral molecular networking of living microbial colonies. Proc. Natl. Acad. Sci. U. S. A. 109, 1743–52 (2012).

15. Protsyuk, I. et al. 3D molecular cartography using LC–MS facilitated by Optimus and ’ili software. Nat. Protoc. 13, 134–154 (2017).

16. Hofmann, A. F. & Hagey, L. R. Key discoveries in bile acid chemistry and biology and their clinical applications: history of the last eight decades. J. Lipid Res. 55, 1553–95 (2014).

17. Yang, J. Y. et al. Molecular networking as a dereplication strategy. J Nat Prod 76, 1686–1699 (2013).

18. Hofmann, A. F. & Eckmann, L. How bile acids confer gut mucosal protection against bacteria. Proc. Natl. Acad. Sci. U. S. A. 103, 4333–4 (2006).

19. Hartmann, A. C. et al. Meta-mass shift chemical profiling of metabolomes from coral reefs. Proc. Natl. Acad. Sci. U. S. A. 114, (2017).

20. Wahlström, A., Sayin, S. I., Marschall, H.-U. & Bäckhed, F. Intestinal Crosstalk between Bile Acids and Microbiota and Its Impact on Host Metabolism. Cell Metab. 24, 41–50 (2016).

21. McDonald, D. et al. American Gut: an Open Platform for Citizen Science Microbiome Research. mSystems 3, e00031–18 (2018).

22. Shalapour, S. et al. Inflammation-induced IgA+ cells dismantle anti-liver cancer immunity. Nature 551, 340–345 (2017).

23. Manor, O. et al. Metagenomic evidence for taxonomic dysbiosis and functional imbalance in the gastrointestinal tracts of children with cystic fibrosis. Sci. Rep. 6, 22493 (2016).

24. Strecker, A. Untersuchung der Ochsengalle. Ann. der Chemie und Pharm. 65, 1–37 (1848).

25. Bernal, J. D. Crystal Structures of Vitamin D and Related Compounds. Nature 129, 277–278 (1932).

26. Gustafsson, B. E., Gustafsson, J. A. & Sjövall, J. Intestinal and fecal sterols in germfree and conventional rats. Bile acids and steroids 172. Acta Chem. Scand. 20, 1827–35 (1966).

27. Midtvedt, T. Microbial bile acid transformation. Am. J. Clin. Nutr. 27, 1341–1347 (1974).

28. Gérard, P. & Philippe. Metabolism of Cholesterol and Bile Acids by the Gut Microbiota. Pathogens 3, 14–24 (2013).

29. Buffie, C. G. et al. Precision microbiome reconstitution restores bile acid mediated resistance to Clostridium difficile. Nature 517, 205–208 (2015).

30. Ma, C. et al. Gut microbiome-mediated bile acid metabolism regulates liver cancer via NKT cells. Science 360, eaan5931 (2018).

31. Sumner, L. W. et al. Proposed minimum reporting standards for chemical analysis Chemical Analysis Working Group (CAWG) Metabolomics Standards Initiative (MSI). Metabolomics 3, 211–221 (2007).

