## Supplementary figures and images for "Chemical Impacts of the Microbiome Across Scales Reveal Novel Conjugated Bile Acids"

### Fig. S1

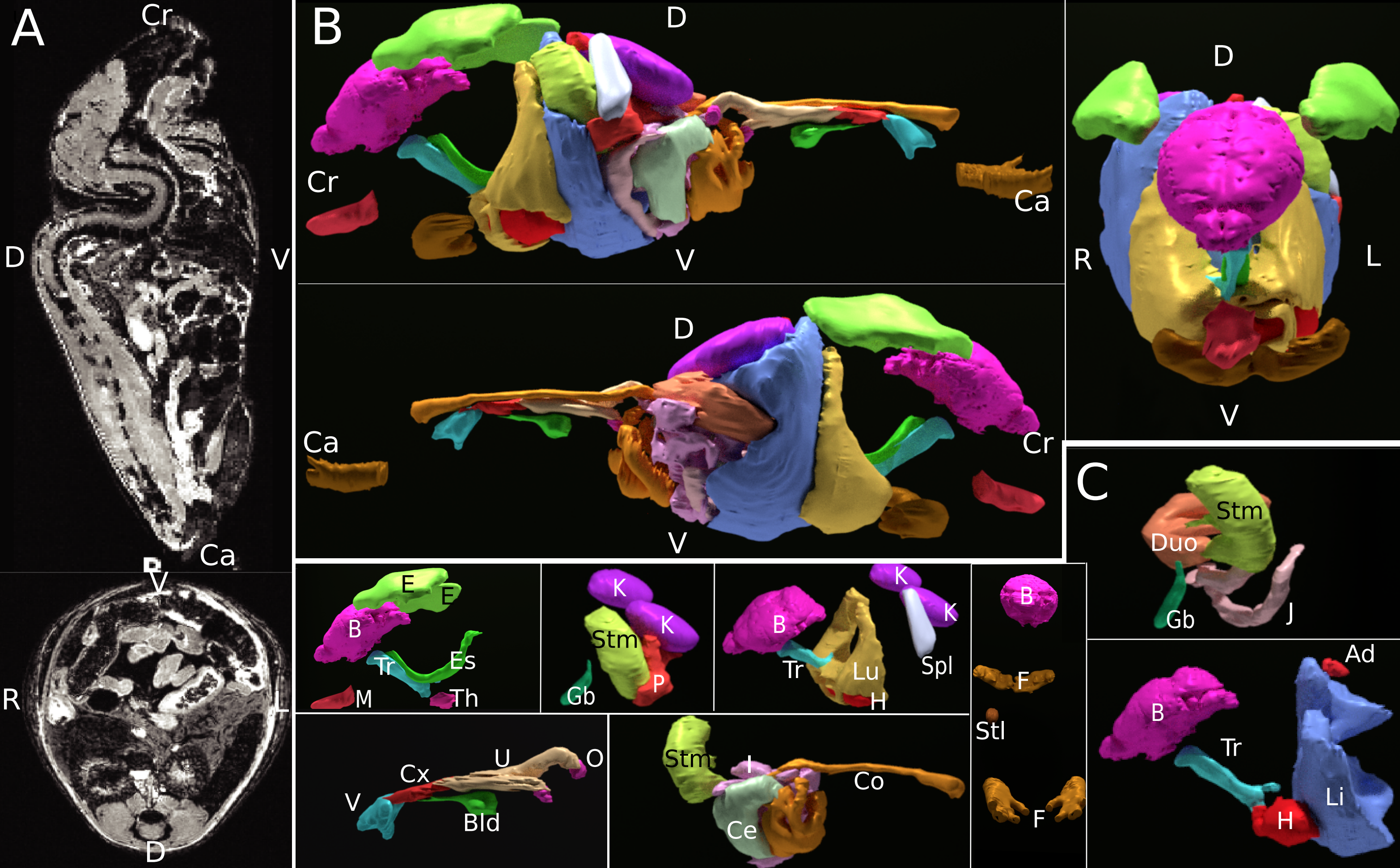

### Fig. S2

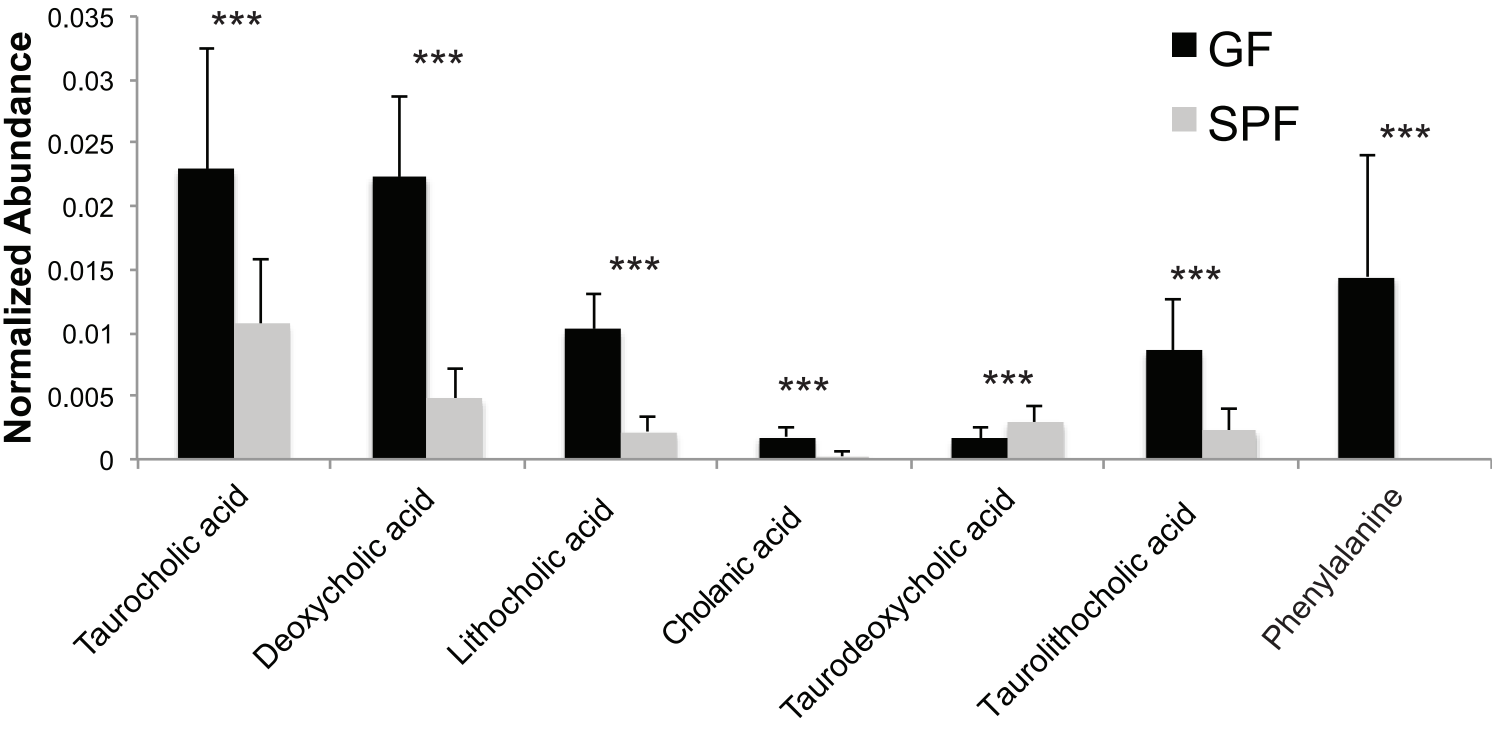

### Fig. S3

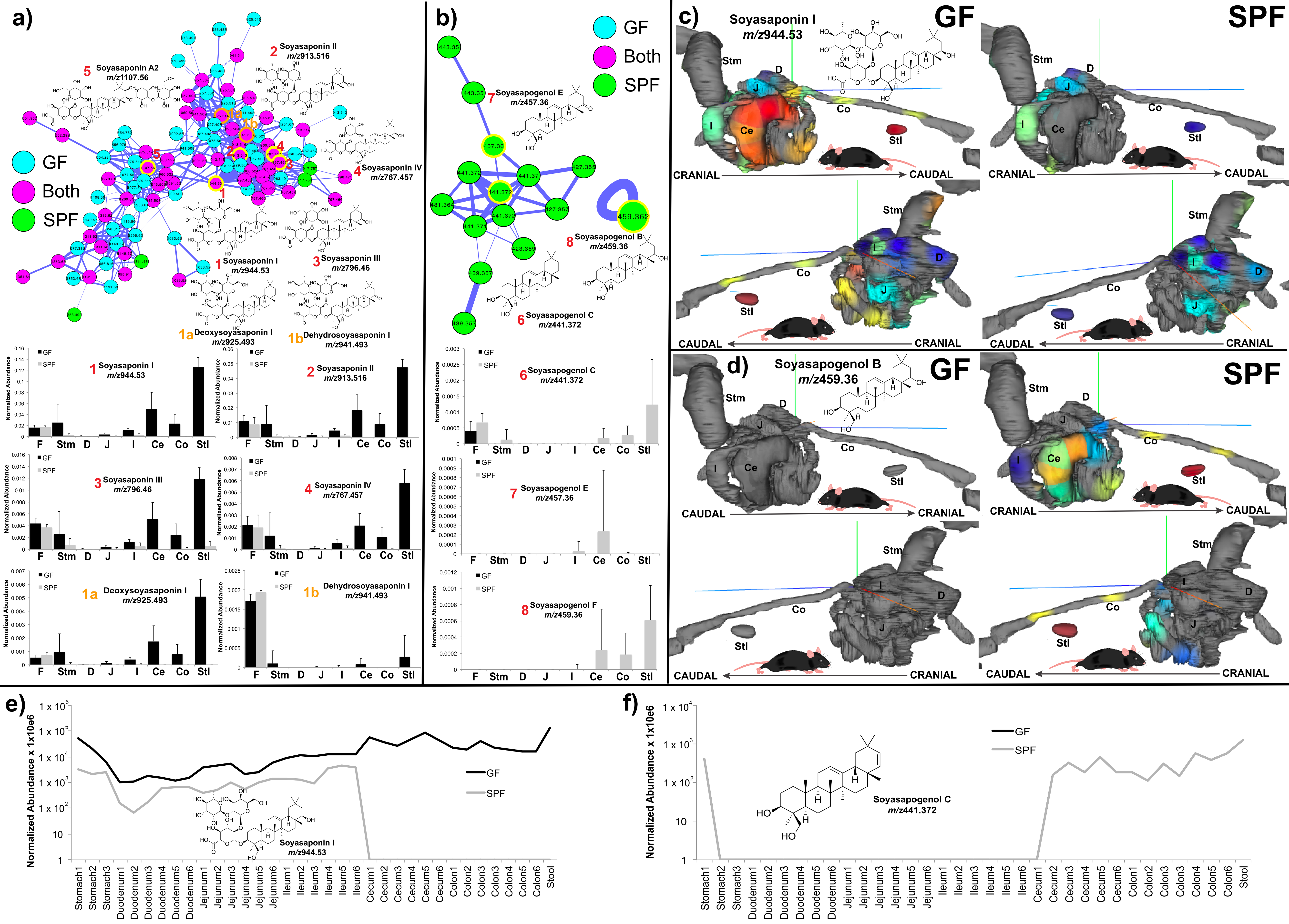

### Fig. S4

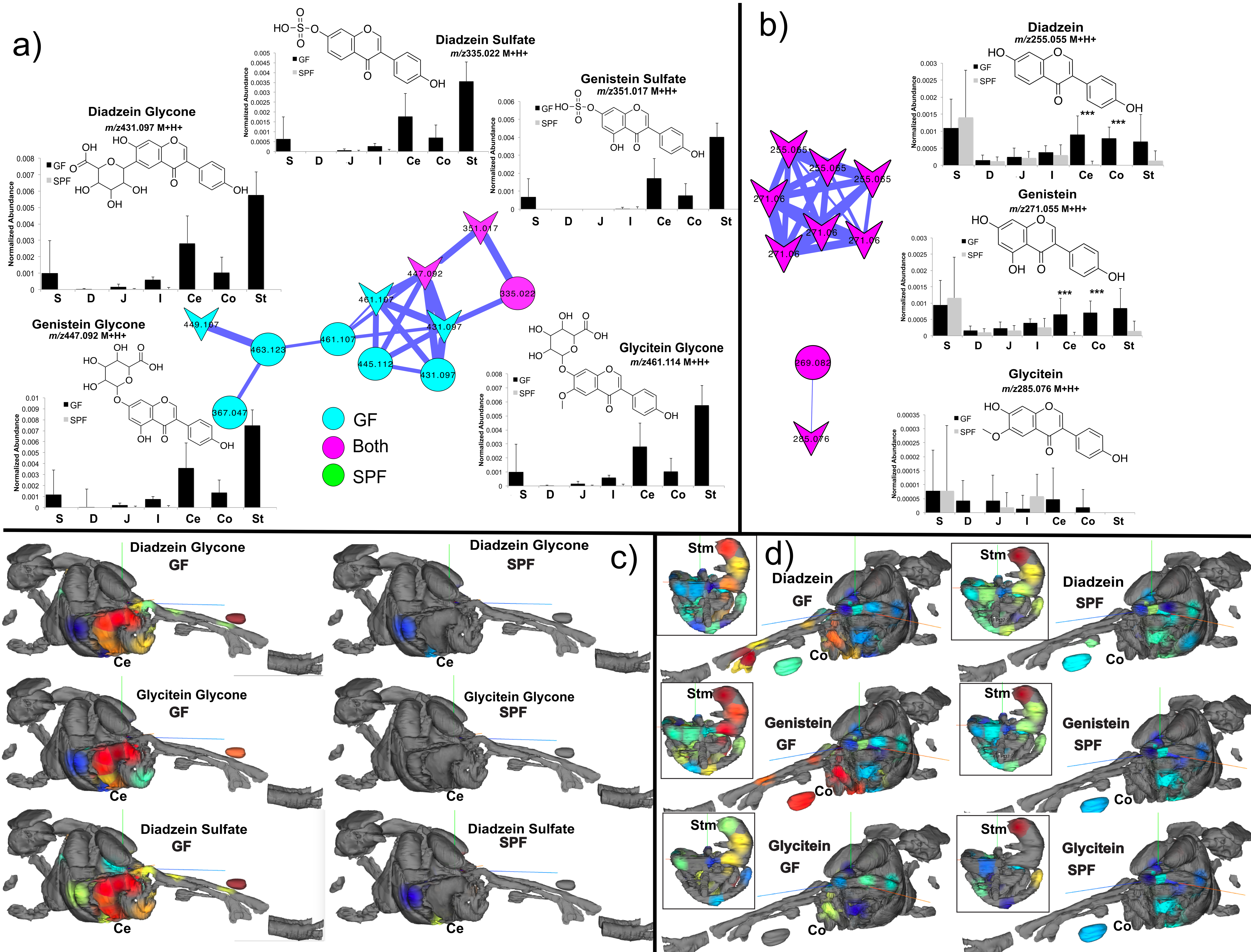

### Fig. S5

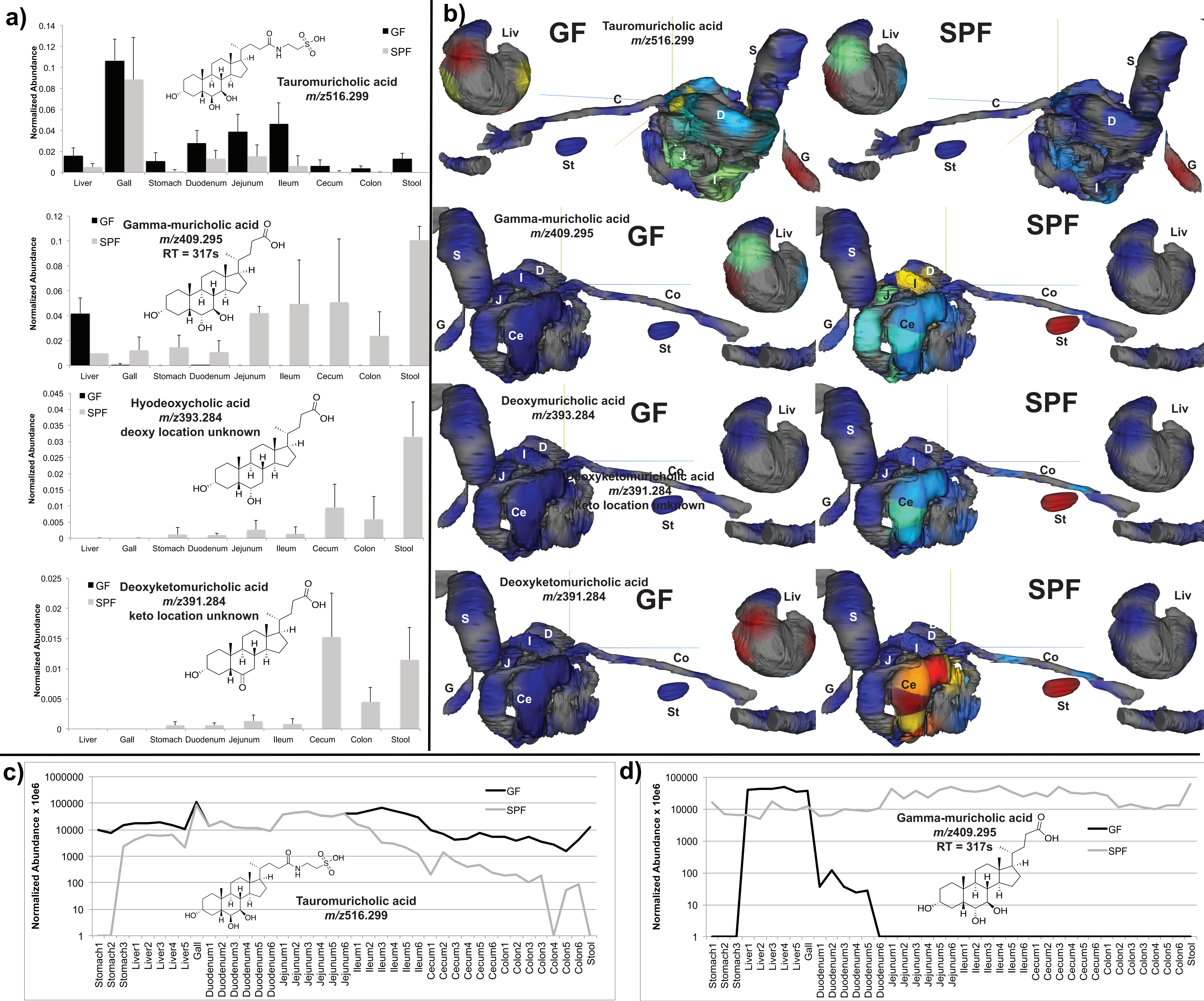

### Fig. S6

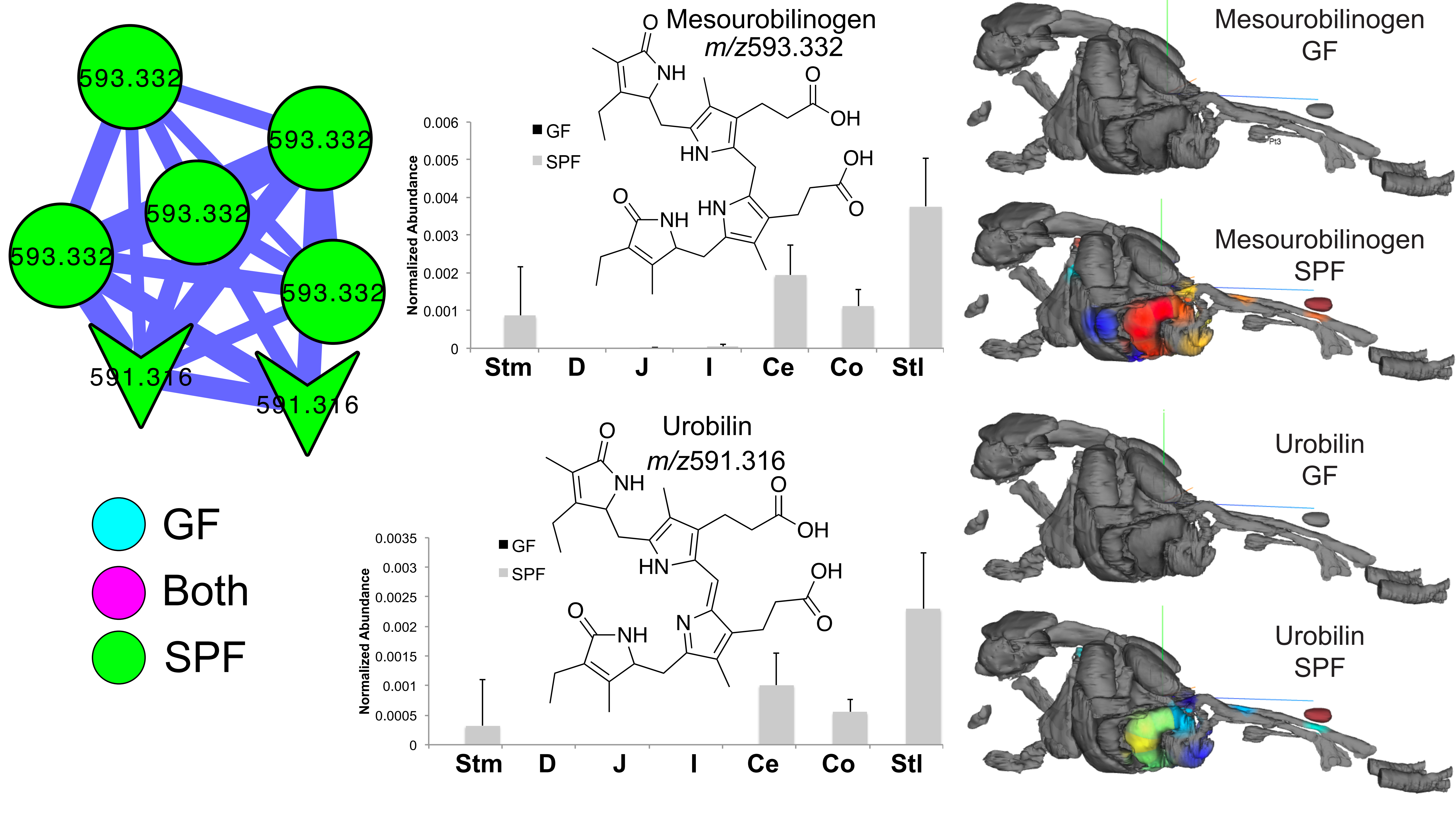

### Fig. S7

#### Amino acid conjugates in colonized murine jejunum.

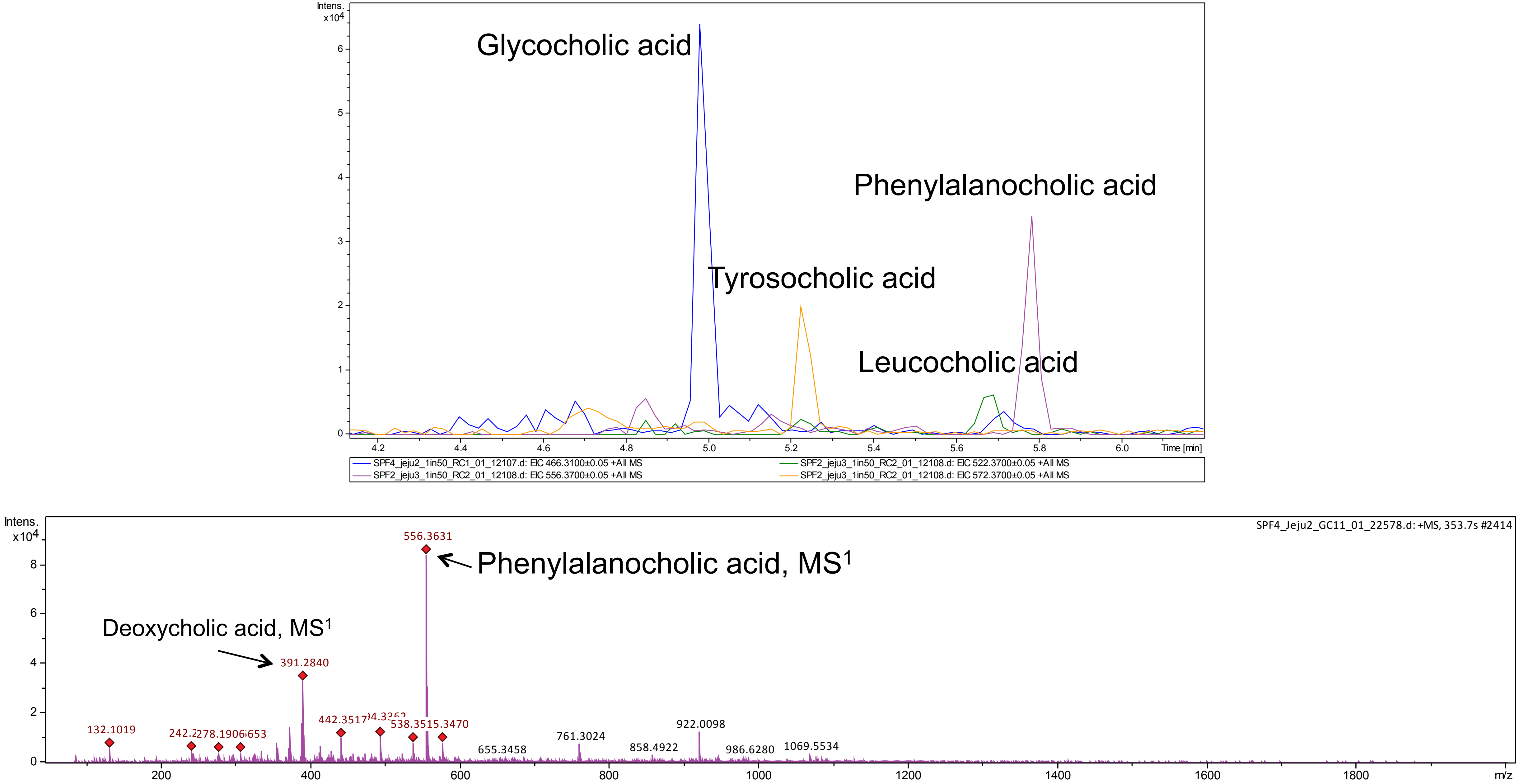

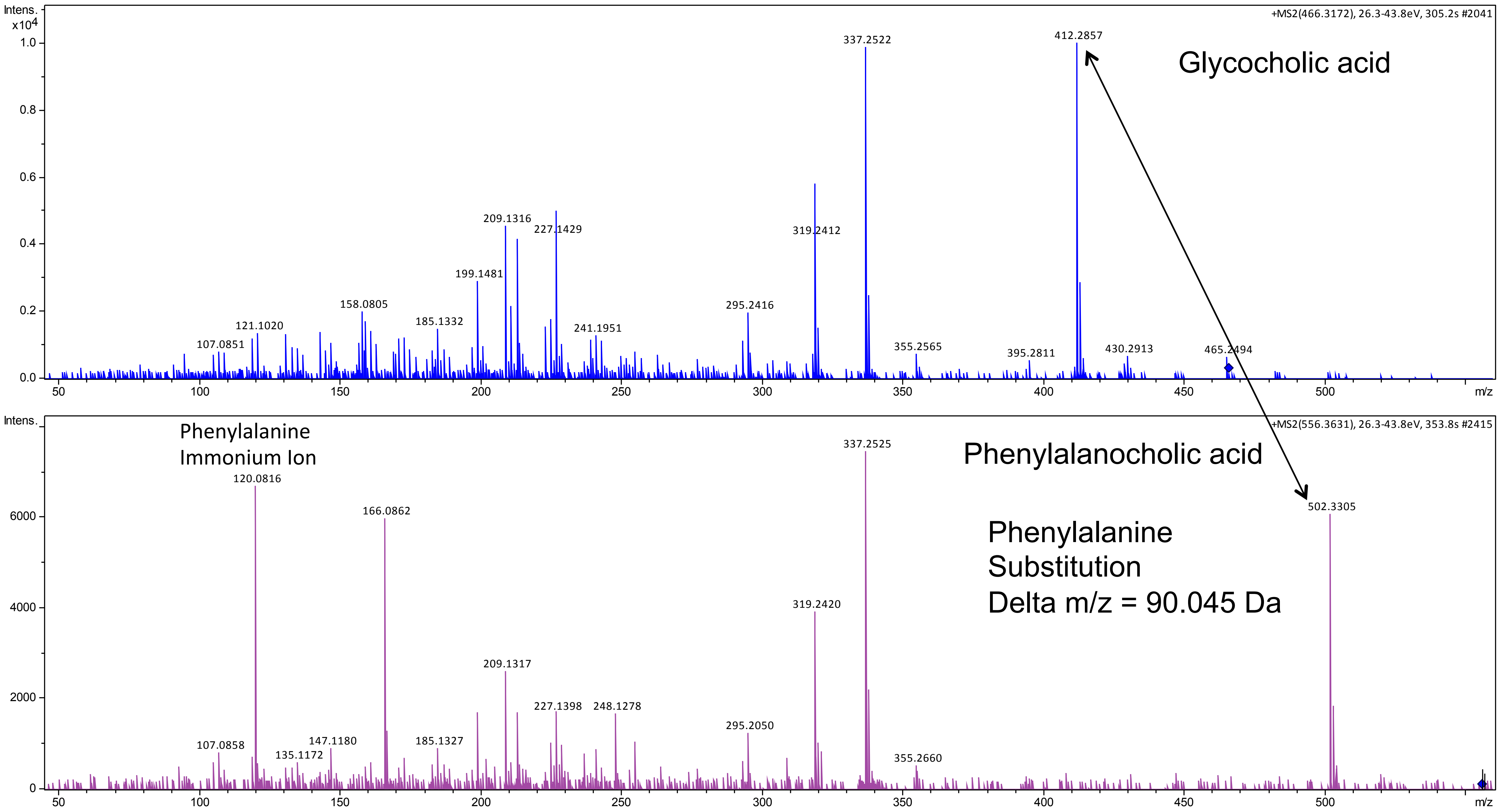

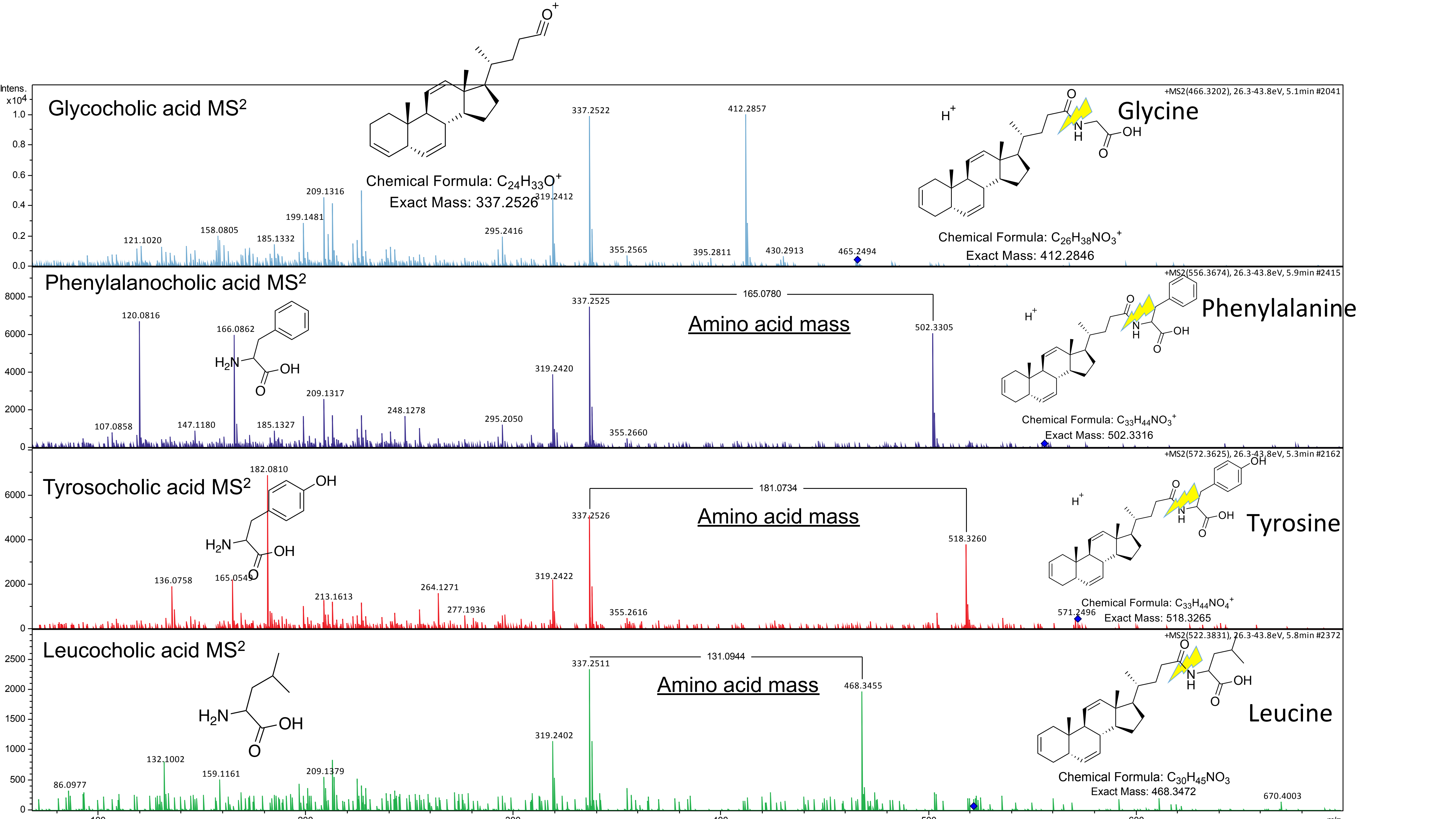

### Fig. S8

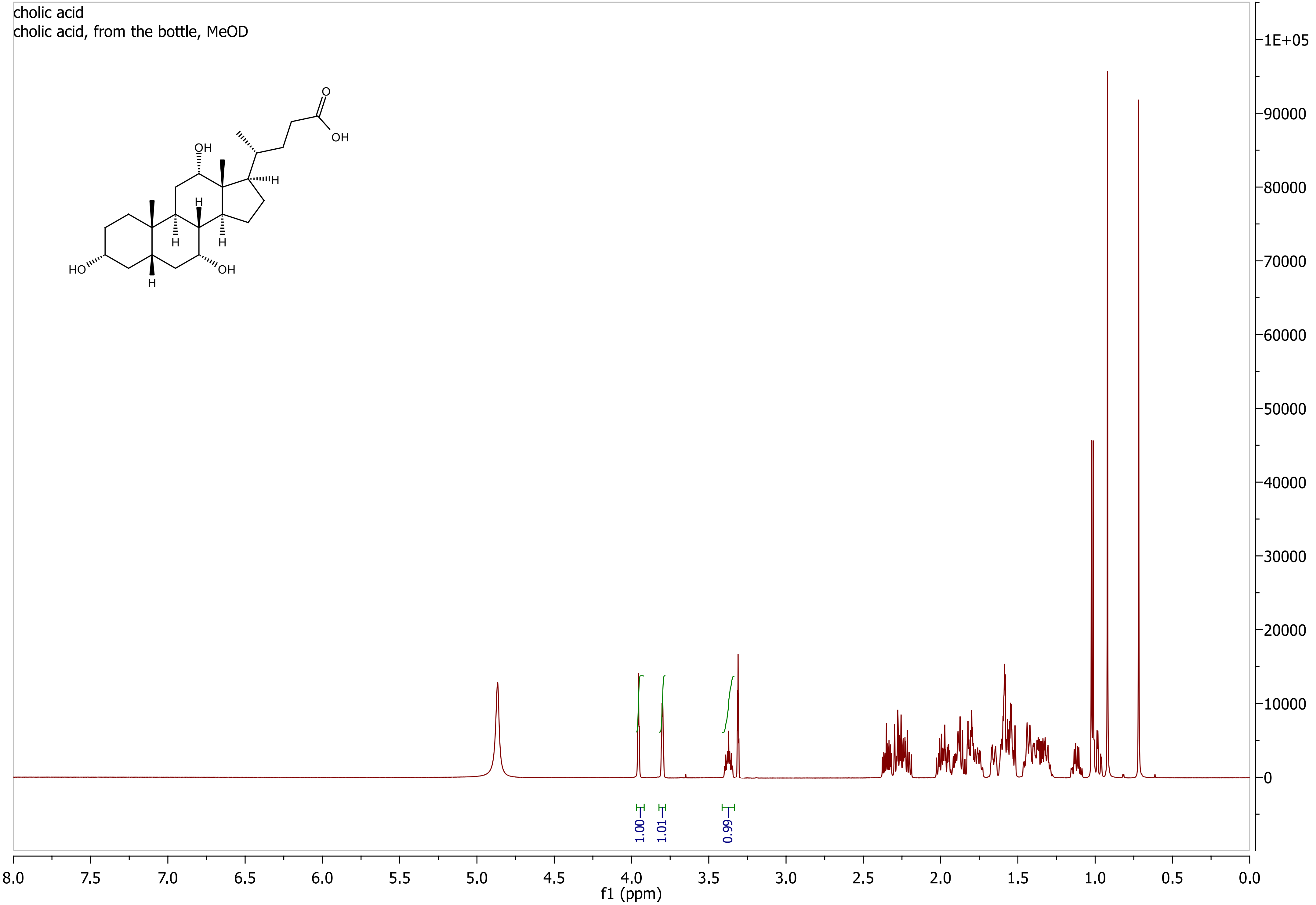

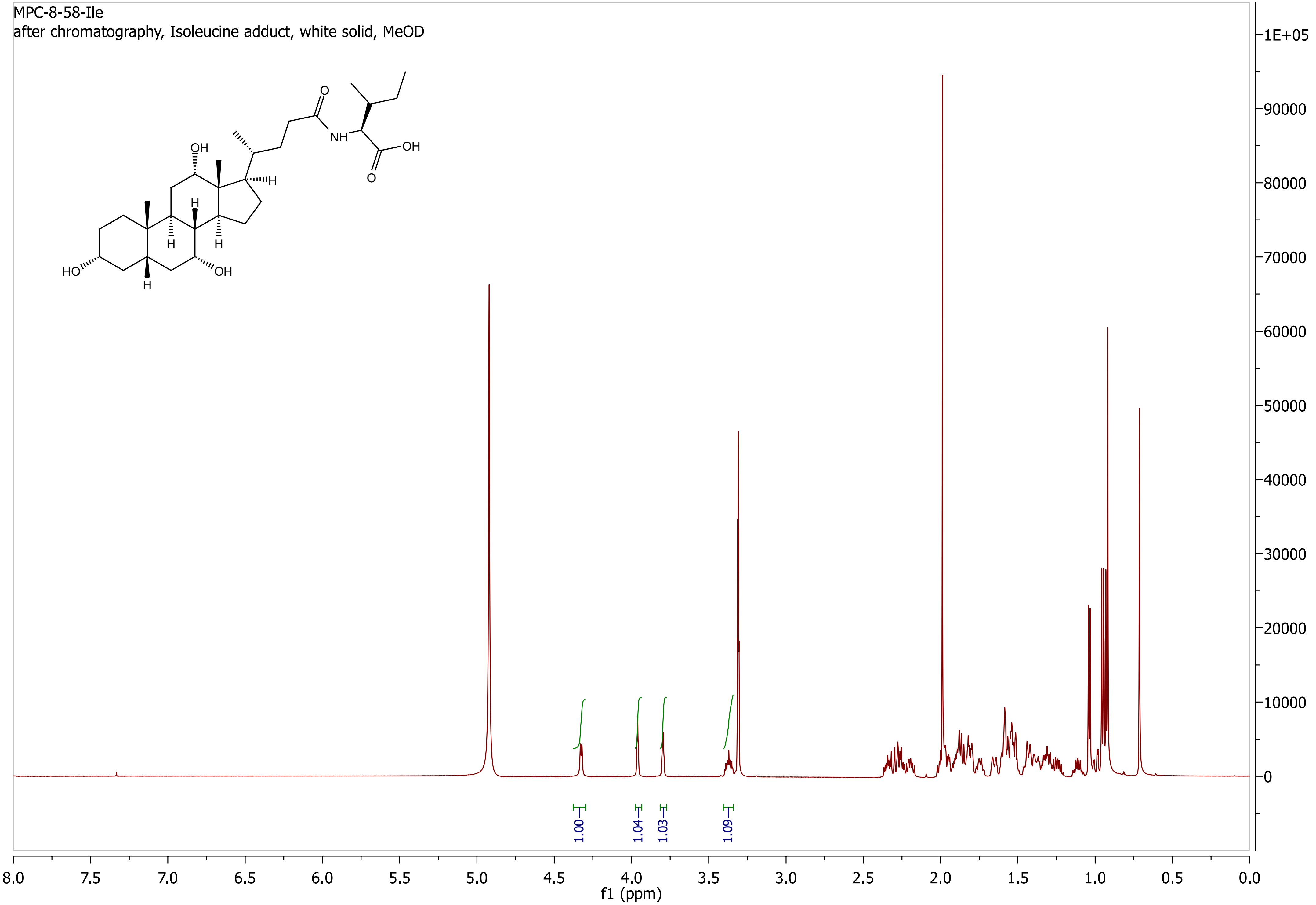

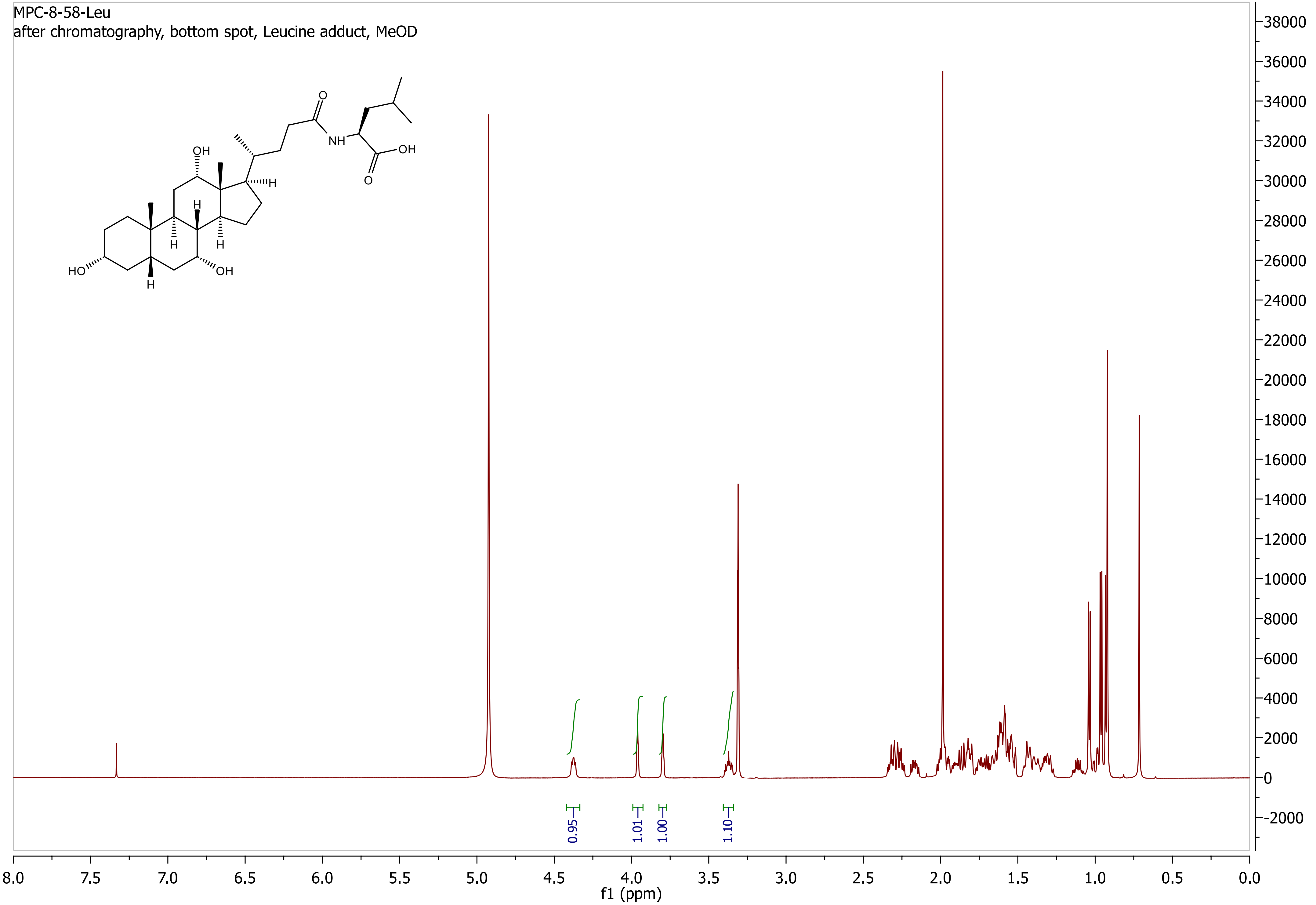

MPC-8-58-Phe
after chromatography, Phenylalanine adduct, white solid, MeOD

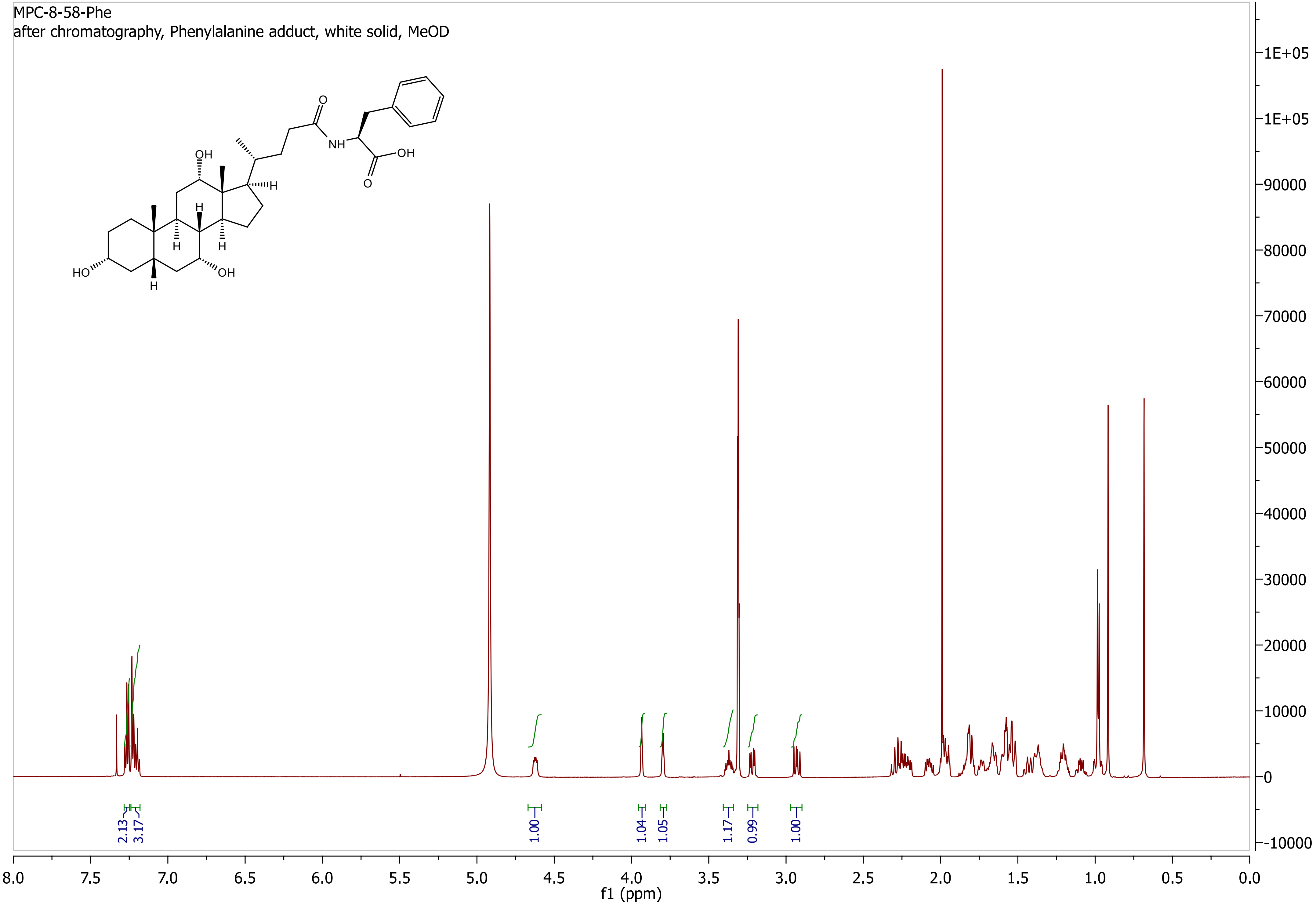

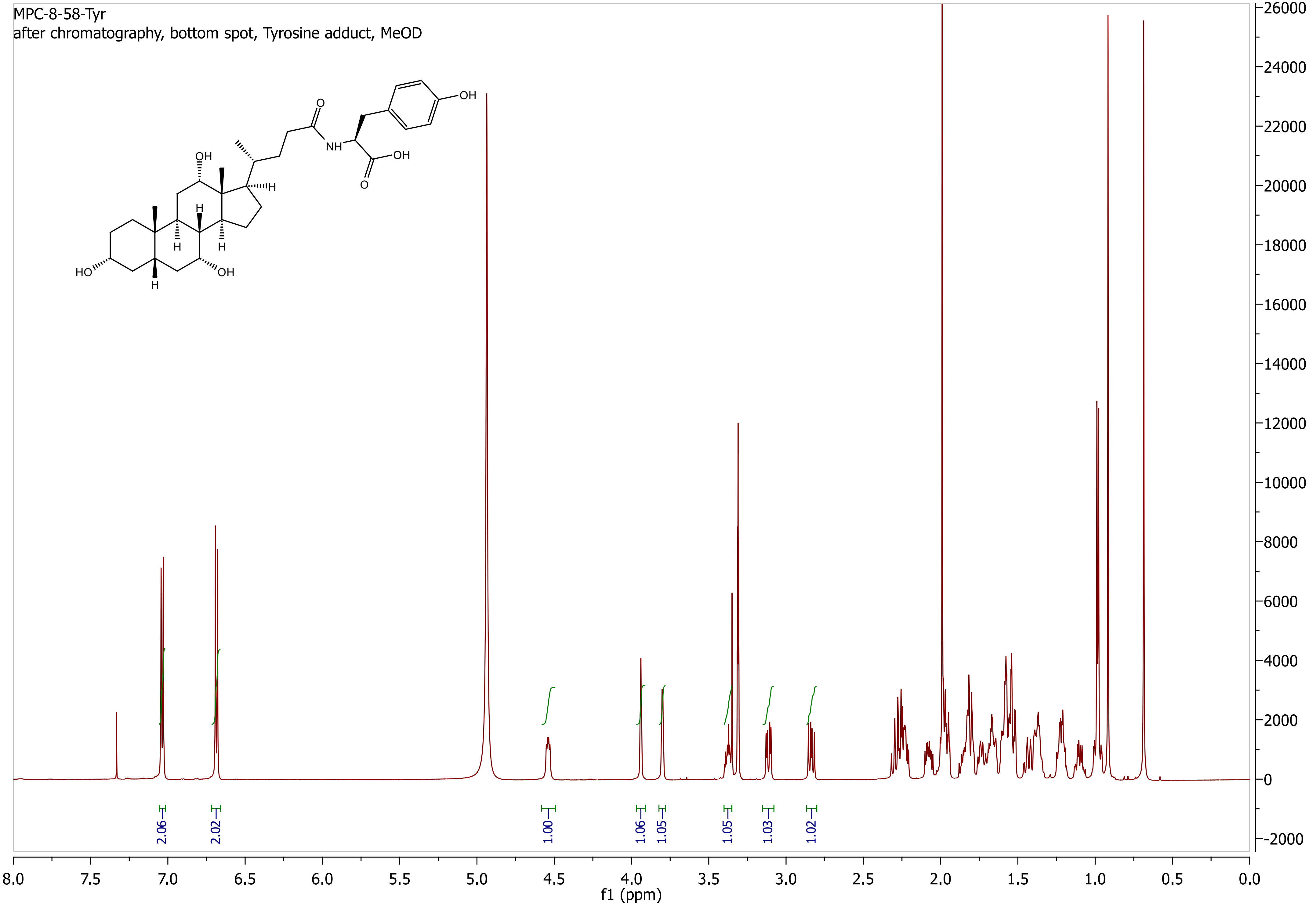

### Fig. S10

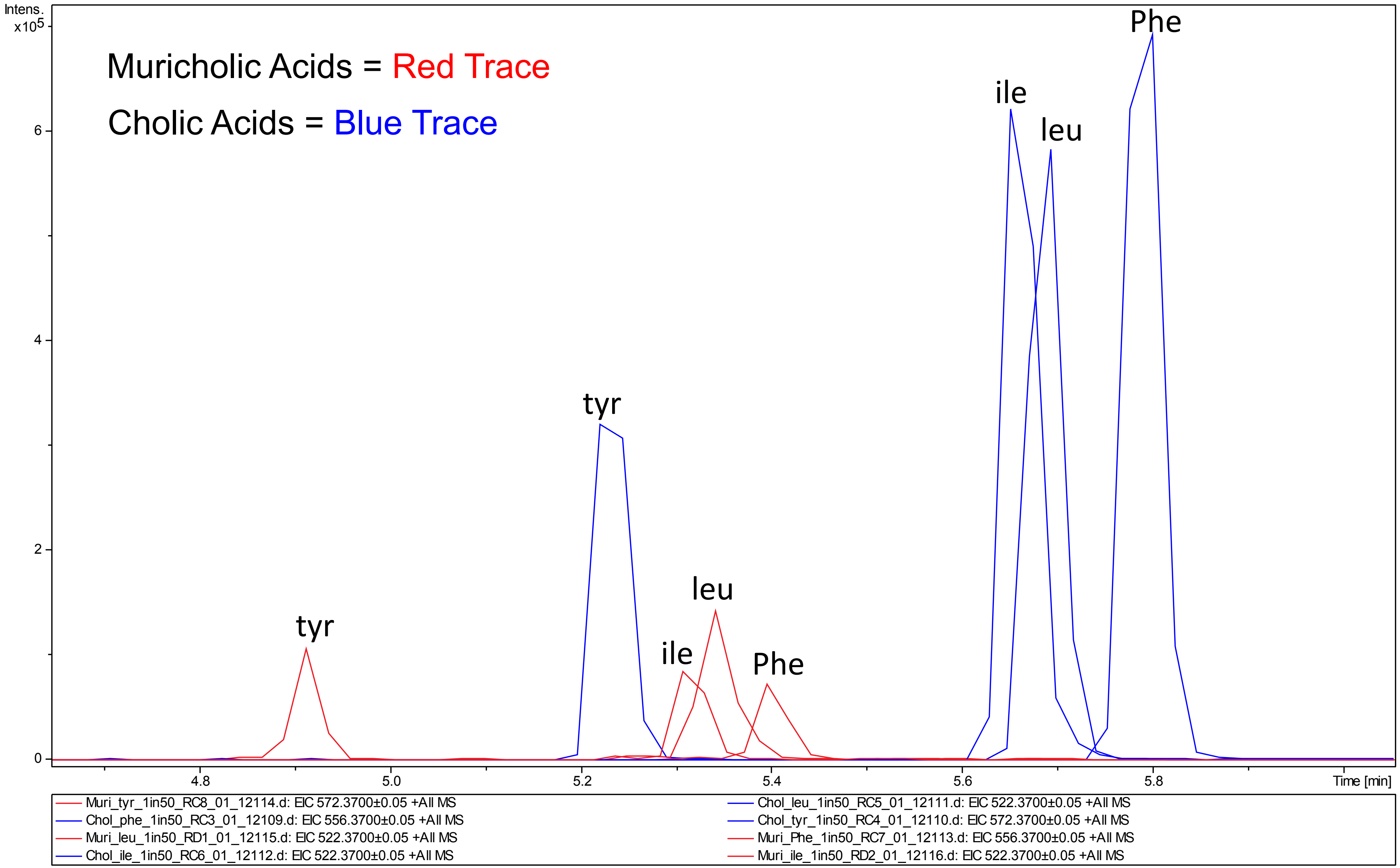

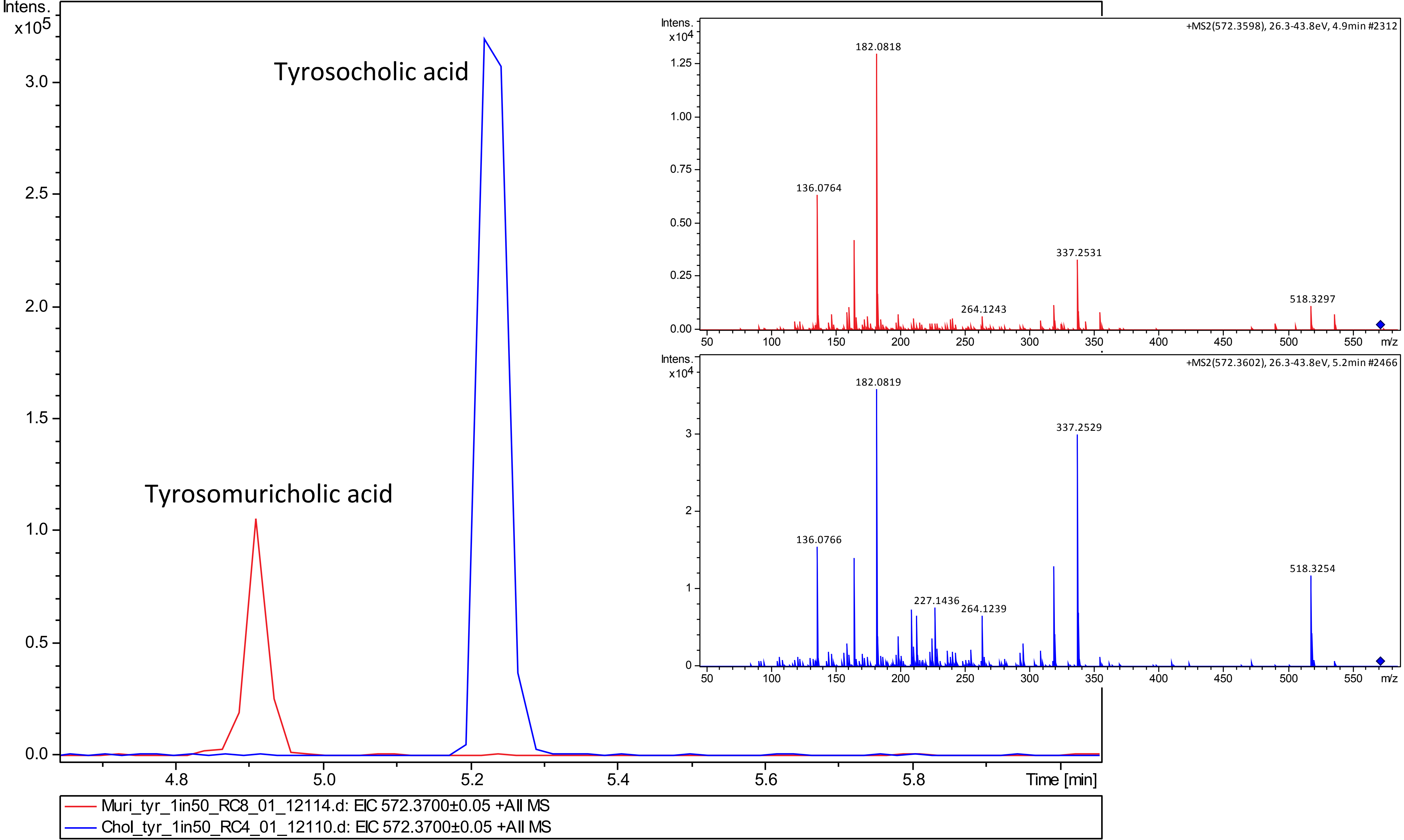

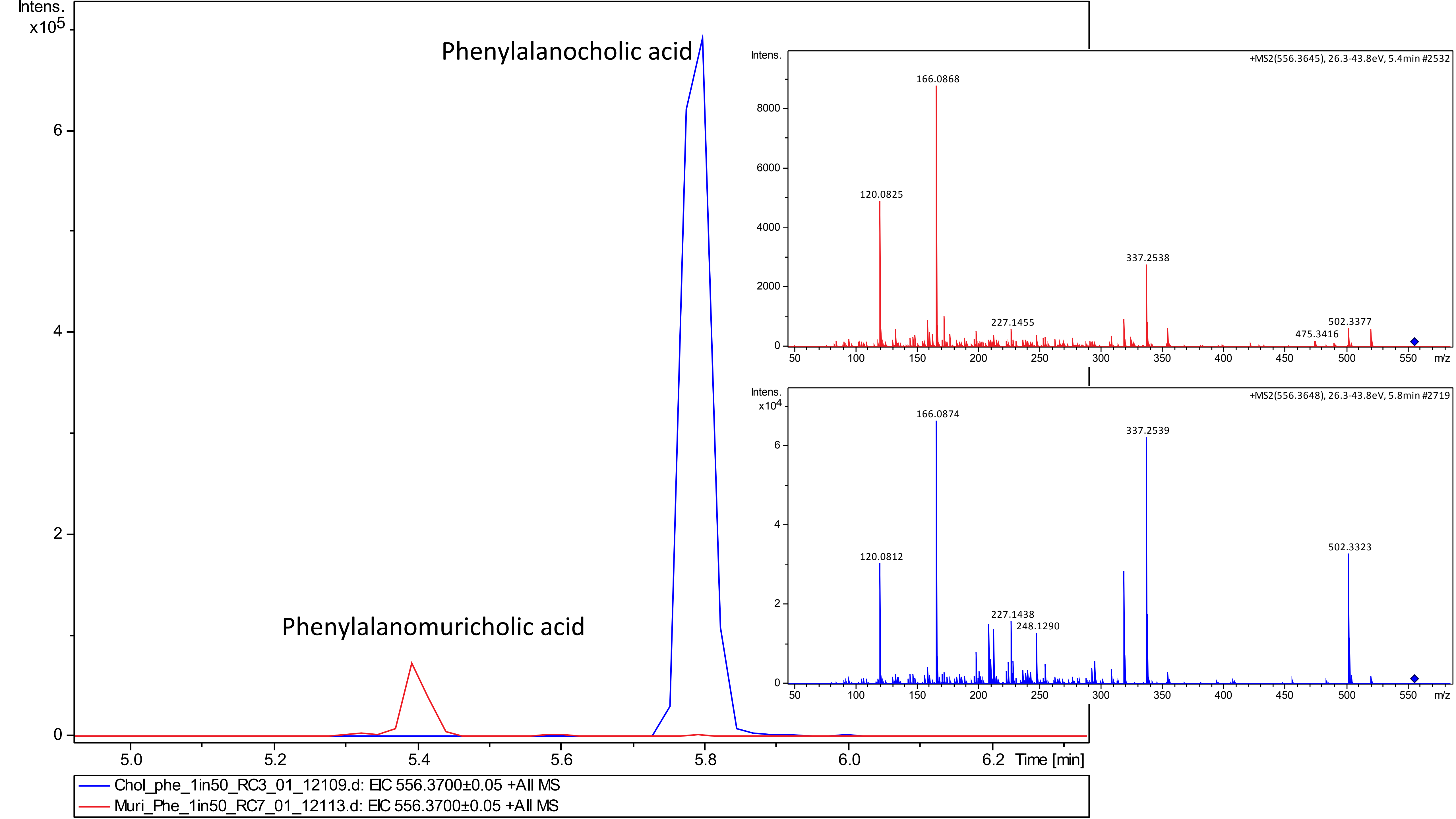

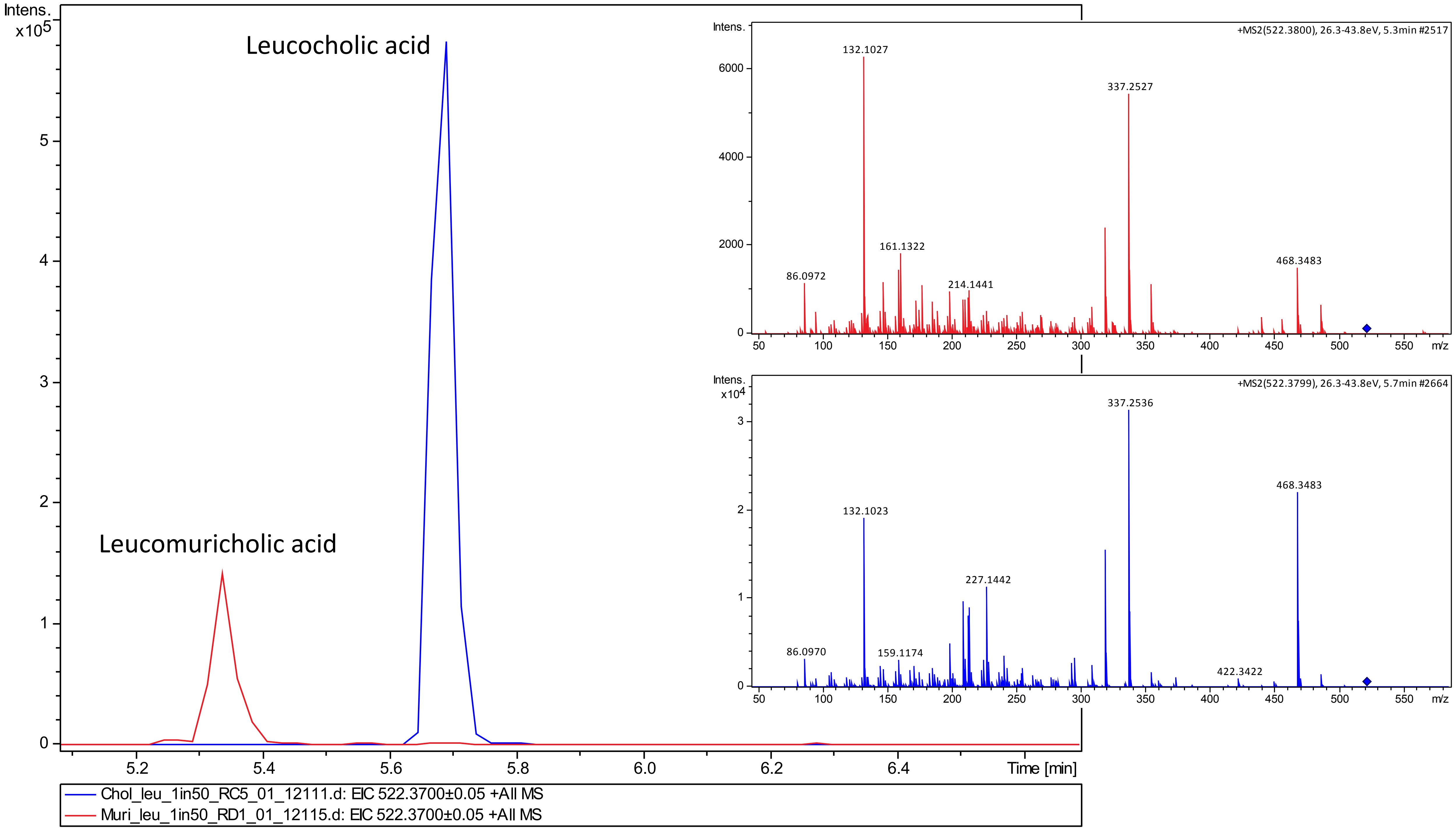

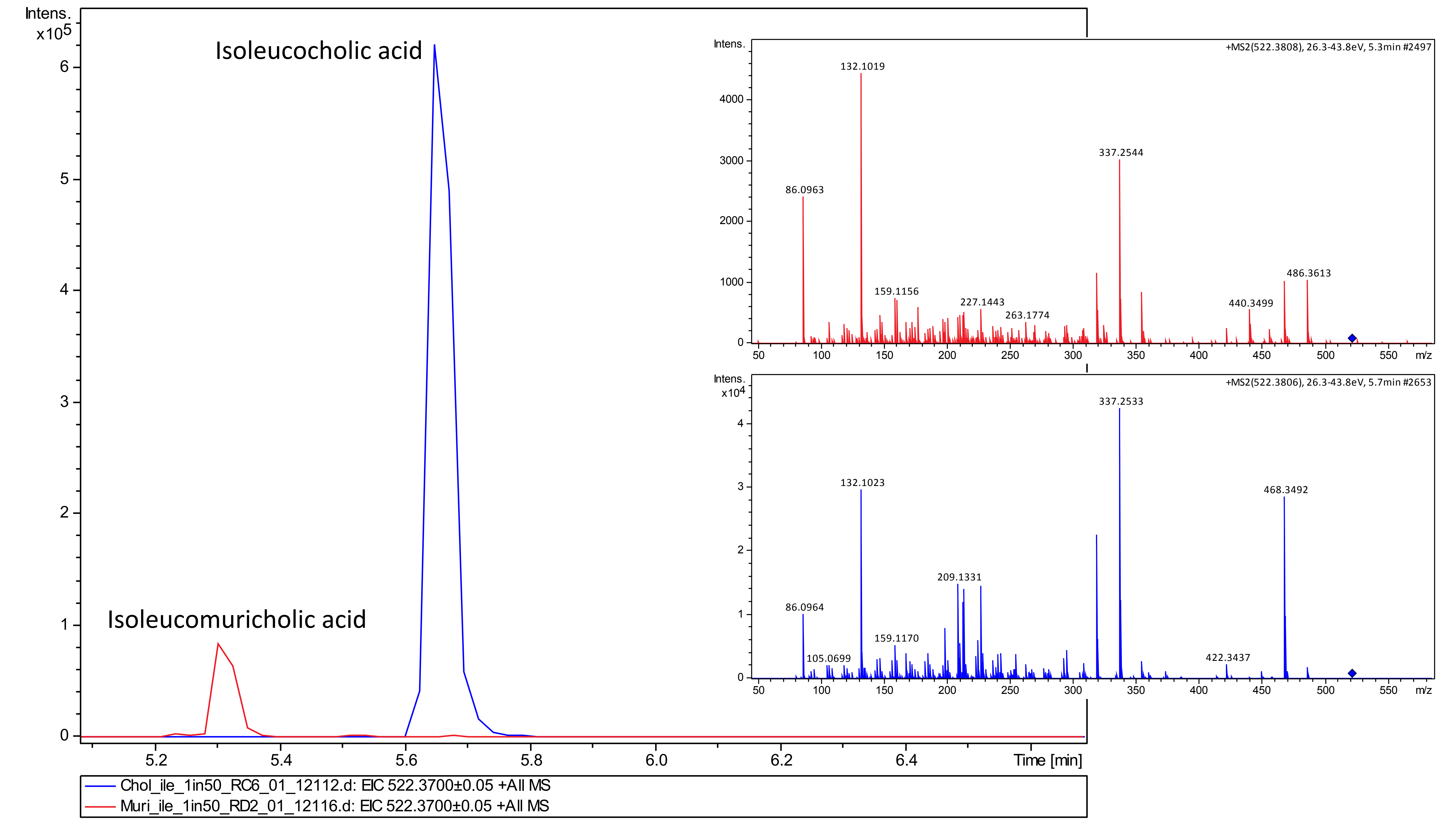

### Fig. S11

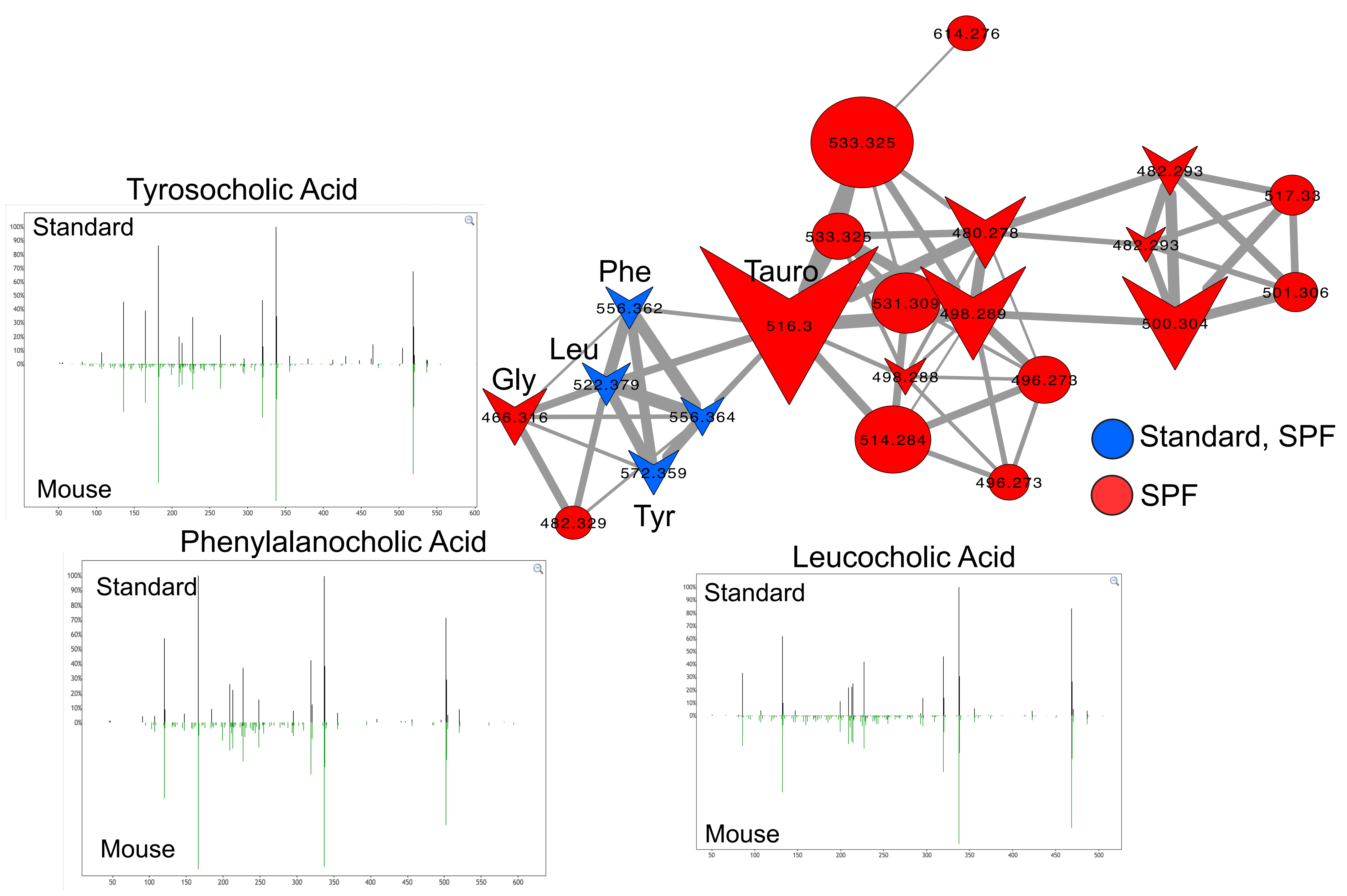

### Fig. S12

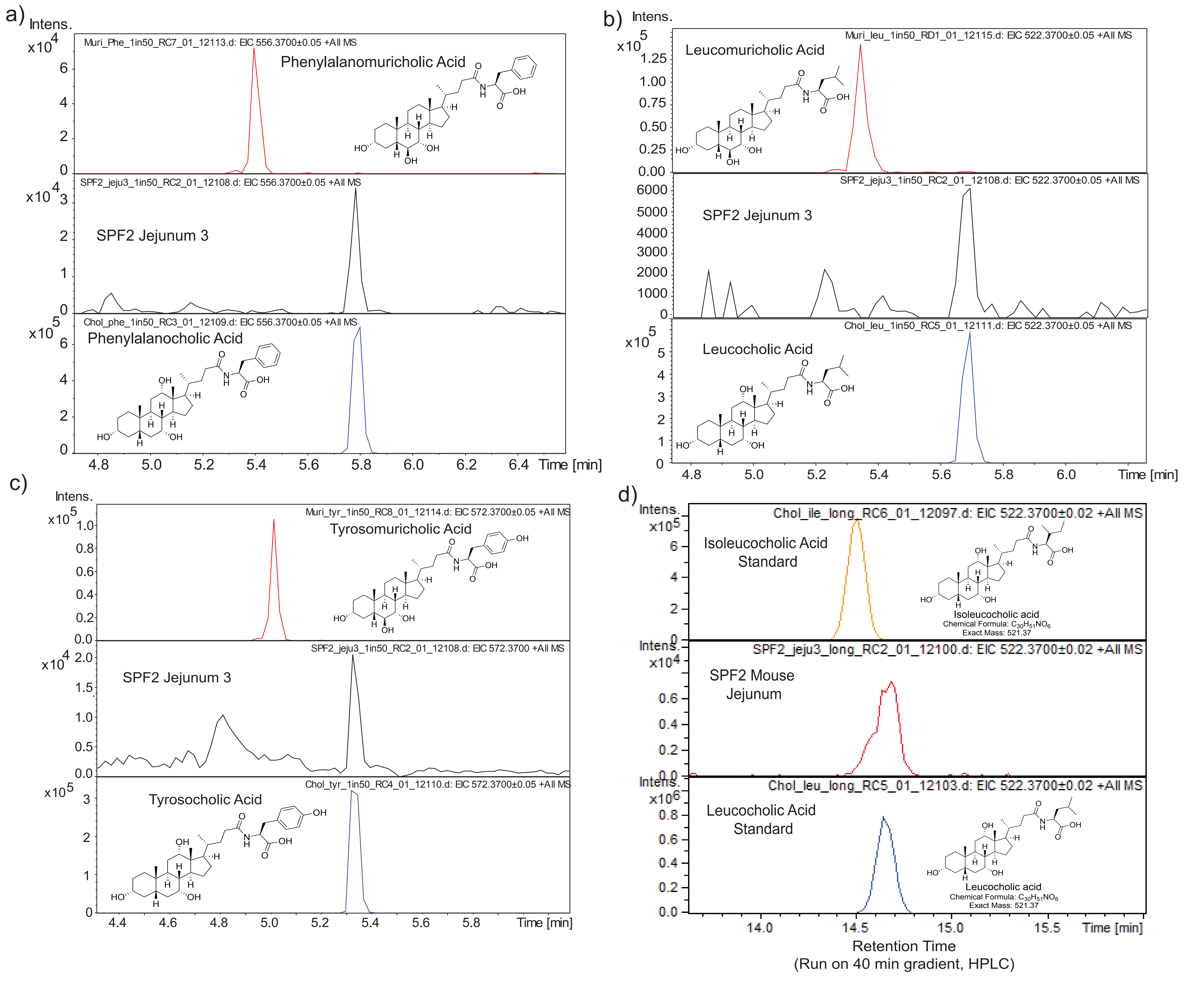

### Fig. S13

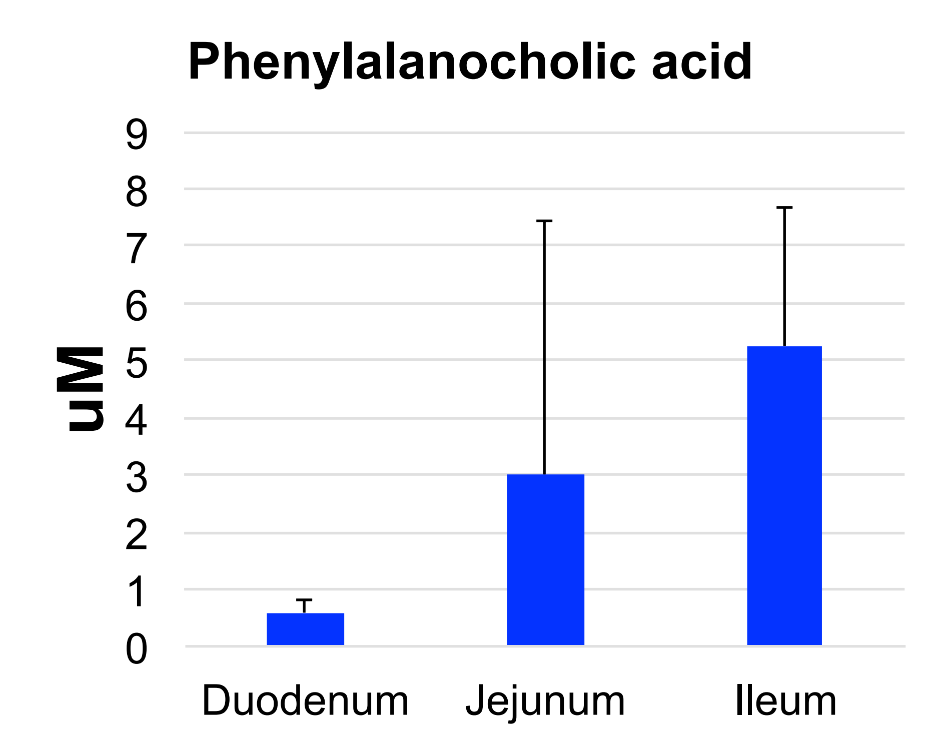

### Fig. S14

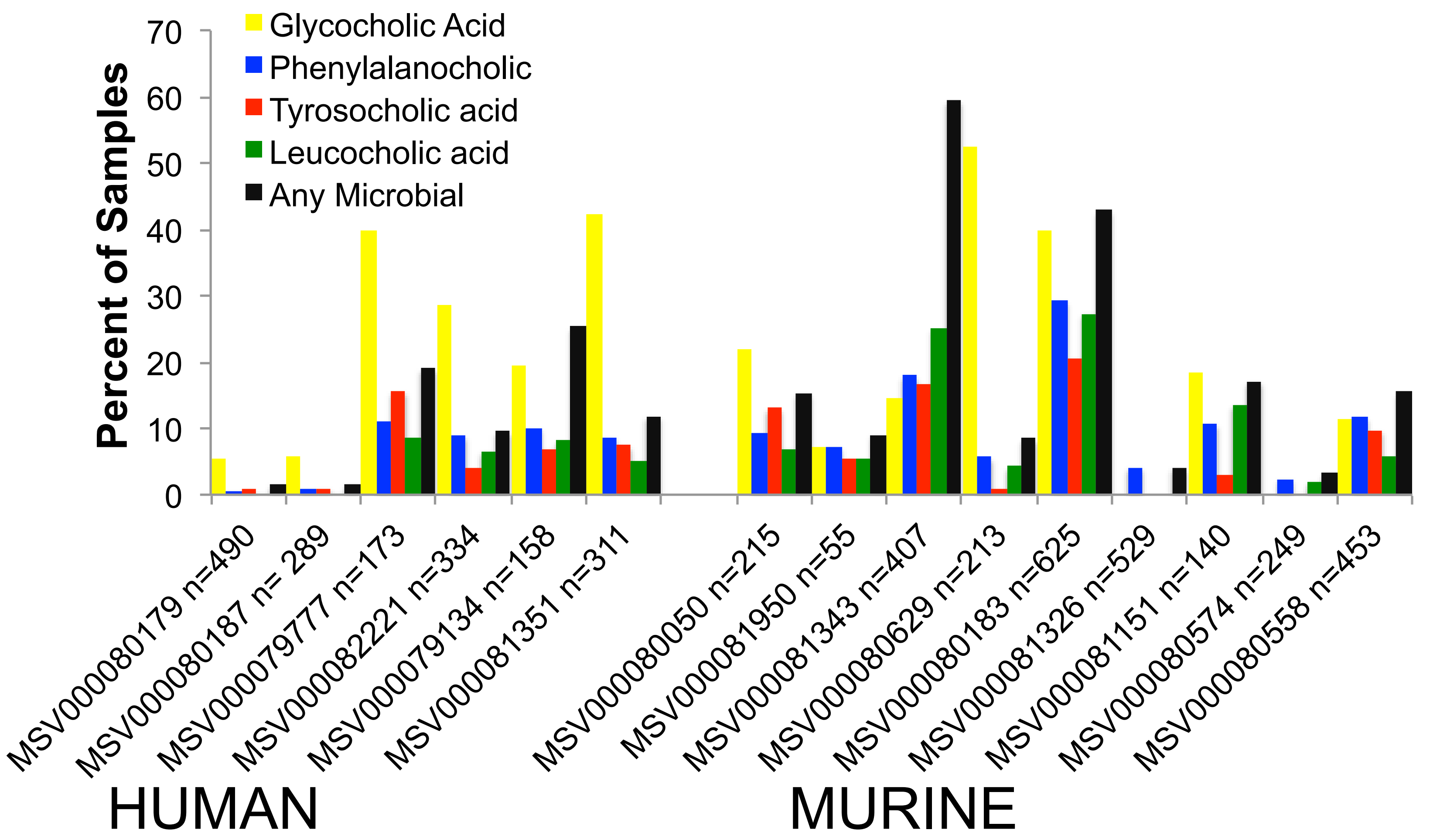

### Fig. S15

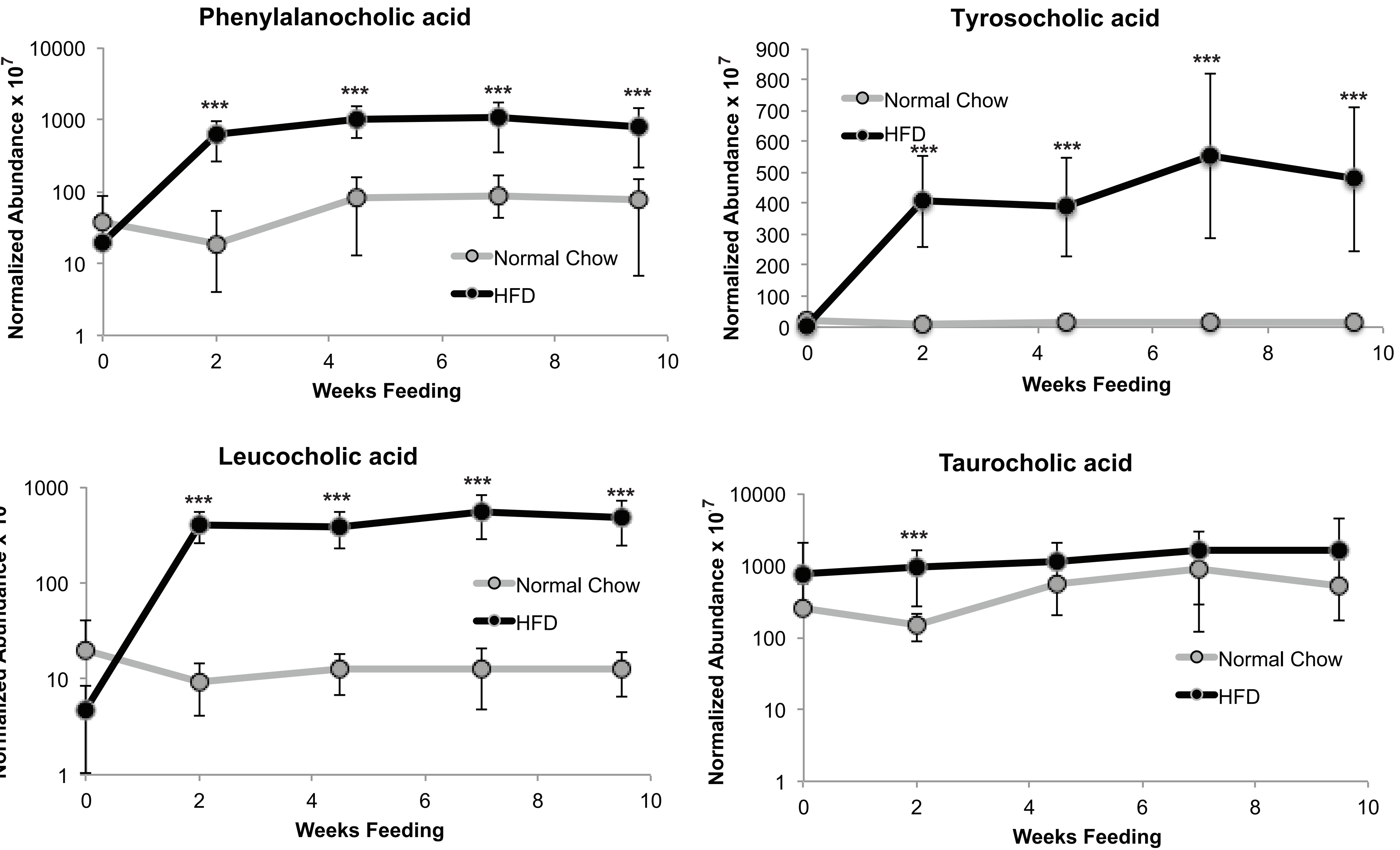
