## Supplementary material for "Chemical Impacts of the Microbiome Across Scales Reveal Novel Conjugated Bile Acids": Fig. S9

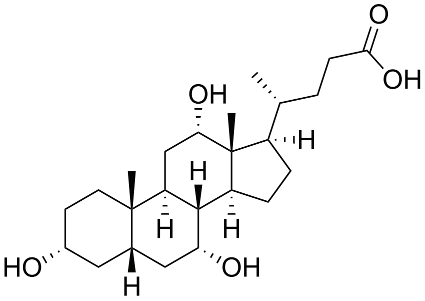

Cholic acid

Chemical Formula: $C_{24}H_{40}O_5$

Exact Mass: 408.29

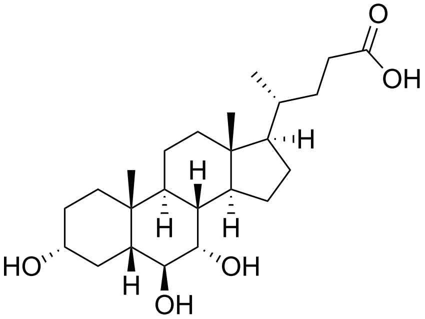

$\alpha$-muricholic acid

Chemical Formula: $C_{24}H_{40}O_5$

Exact Mass: 408.29

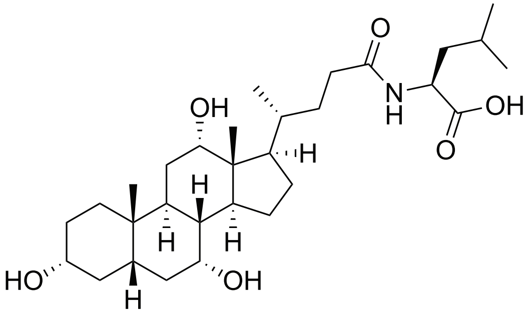

Leucocholic acid

Chemical Formula: $C_{30}H_{51}NO_6$

Exact Mass: 521.37

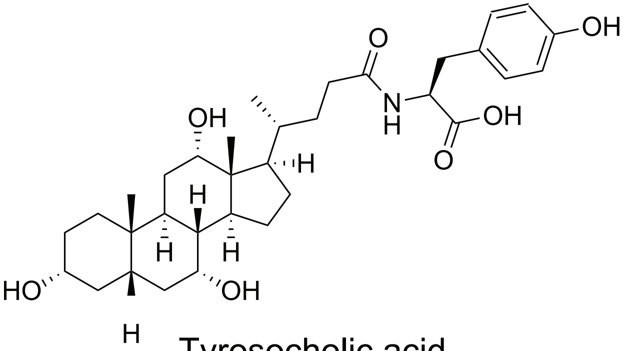

Tyrosocholic acid

Chemical Formula: $C_{33}H_{49}NO_7$

Exact Mass: 571.35

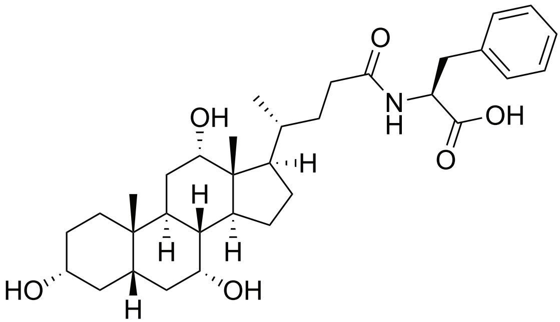

Phenylalanocholic acid

Chemical Formula: $C_{33}H_{49}NO_6$

Exact Mass: 555.36
