## Supplemental Data for "Chemical Impacts of the Microbiome Across Scales Reveal Novel Conjugated Bile Acids"

**Supplemental Figure Legends**

Figure S1. Schematic illustrating the generation of the 3-D mouse model for mapping multi-omics data in this study. a) A high resolution MRI of a representative laboratory mouse was used to trace organs in each stack of the MRI from cranial/caudal, dorsal/ventral and left/right planes. b) The complete model from side and frontal views. Each organ was traced using the Invesalius software in each plane and a 3D rendering of the organ was produced one-by-one to build the entire murine model. c) Visualization of each organ in the model by removing overlapping sections. E-ear, Es-esophagus, B-brain, M-mouth, Tr-trachea, Th-thymus, H-heart, Ad-adrenal gland, Li-liver, Lu-lung, Spl-spleen, K-kidney, F-feet, Stl-stool, D-duodenum, G-gall bladder, J-jejunum, Stm-stomach, P-pancreas, I-ileum, Ce-cecum, Co-colon, O-ovary, U-uterus, Cx-cervix, V-vagina

Figure. S2. Normalized abundance of metabolites differentiating GF and SPF mouse liver samples. All liver samples were classified as either GF or SPF and a random forests classification was used to identify variables of importance. Significance was then tested with the Wilcoxon Rank-Sum test. Known metabolites identified are shown and constituted particular microbially influenced bile acids and phenylalanine. ***=p<0.001

Figure S3. a) Molecular network cluster of soyasaponins colored by source of each node as GF, SPF or shared. Structures of corresponding molecules are shown in nodes highlighted in yellow according to the numbering scheme. Mean abundance and standard deviations of the means of each soyasaponin metabolite from the murine GI tract in the GF and SPF mice and the food pellet (F=food, Stm=Stomach, D=Duodenum, J=Jejunum, I=Ileum, Ce=Cecum, Co=Colon, Stl=Stool). b) Molecular family of soyasapogenols, their structures and relative abundances in GF and SPF gut organs. c) ‘*ili* 3-D model visualization of the normalized abundance of soyasaponin I in the murine GI tract. Abundance of the metabolite is indicated according to the viridis spectrum (high red/hot colors, low blue/cool colors). d) ‘*ili* 3D cartography of the normalized abundance of soyasapogenol B onto an MRI organ model of the mice. e) Mean normalized abundance of soyasaponin I through all GI sample locations in the GF and SPF mice. f) Mean normalized abundance of soyasapogenol through all GI sample locations.

Figure S4. Microbial metabolism of plant isoflavones in GF and SPF metabolomics data. a) Structures, molecular network and normalized abundance of glycone isoflavanoids in the murine GI tract. Nodes are colored according to their source in GF or SPF mice and known library hits are shaped as arrowheads. d) Same information for the aglycones. c) 3D-molecular cartography mapping the abundance of the diadzein and glycitein glycone and suflated forms through entire 3D-mouse model. The normalized abundance of a particular molecule is indicated as a heat map with red being most abundant and blue being lowest abundance. d) 3D-molecular cartography mapping the abundance of the diadzein and glycitein aglycone forms through entire 3D-mouse model. GI tract model is inset for reference.

Figure S5. Microbial metabolism of known bile acids in GF and SPF metabolomics data. a) Normalized abundance of taurocholic acid and secondary bile acids in GF and SPF mice GI tract samples. b) 3D-molecular cartography mapping the abundance of the same bile acids through the mouse GI tract model including liver separated for better visualization. The normalized abundance of a particular molecule is indicated as a heat map with red being most abundant and blue being lowest abundance. c) Normalized abundance of taurocholic acid through the GI tract of GF and SPF mice. d) Normalized abundance of gamma-muricholic acid through the GI tract of GF and SPF mice.

Figure S6. Urobilin and mesourobilinogin structures, molecular network, normalized abundance in GI tract and 3D-molecular cartography through entire murine model.

Figure S7. Chromatography and MS^2^ spectra of novel amino acid conjugated bile acids. Loss of the unique amino acids are shown in the MS^2^ spectrum, which appear as whole amino acids in the lower m/z range.

Figure S8. Nuclear magnetic resonance spectra of synthesized glycocholic acid, phenylalanocholic acid, leucocholic acid, isoleucocholic acid and tyrosocholic acid.

Figure S9. Structures and exact masses of known and novel conjugated bile acids in this study from murine and human gut samples.

Figure S10. LC-MS/MS data of synthetic muricholic acid and cholic acid conjugates. Retention time differences between all conjugates and their subsequent MS^2^ data are shown.

Figure S11. Molecular network of SPF duodenum and synthesized amino acid conjugated bile acids. LC-MS/MS data from synthetic standards was networked with murine samples and spectral matching through molecular networking is indicated by node coloring. Mirror plots showing the alignment between the murine and standards are shown.

Fig. S12. Retention time alignments of novel synthetic muricholic and cholic acid conjugates with those found in a murine jejunum sample. a) Phenylalanocholic acid, b) leucocholic acid, c) tyrosocholic acid. d) Retention time analysis of synthetic isoleucocholic acid, leucocholic acid and a murine jejunum sample run on a long gradient HPLC column to separate isomeric ile/leu conjugates and compare to that detected *in vivo*.

Fig. S13. Mean and standard deviations of amino acid conjugate concentrations found in the murine GI tract according to that of a standard curve created by spiking synthetic standards into a GF murine GI sample.

Figure S14. Proportion of samples where phenylalanocholic acid, tyrosocholic acid and leucocholic acid were found through single spectrum searching of publically available data on GNPS. Massive data set ID’s are shown for each dataset and they are divided as either murine or human GI samples.

Figure S15. Mean normalized abundance of the three novel conjugated bile acids compared to taurocholic acid in mice (ApoE knockout on a C57BL/6J background) fed either HFD or normal chow for 10 weeks. Fecal samples were collected and extracted in 50:50 methanol water and analyzed with LC-MS/MS metabolomics according to^1^. Standard deviations around the means are shown and significance between HFD and normal chow at each time point is tested with the student’s t-test (***=p<0.001).

**Supplemental Results**

**Microbiome Changes through the Murine GI tract.** The 16S rRNA gene microbiome profiles of the GI tract were dominated by Bacteroidales clade S24-7, Firmicutes, *Lactobacillus* and *Akkermansia muciniphila* (Fig. 3b,c). The esophagus, stomach and duodenum had relatively similar profiles, but a dramatic shift in the jejunum with the expansion of *Lactobacillus* and *A. muciniphila* and a decrease in the relative abundance of Bacteroidales S24-7 was evident. The community transitioned again through the ileum with a further expansion of *Lactobacillus*. At the cecum an abrupt transition was observed with a reduction of *Lactobacillus* and increase in the relative abundance of Firmicutes (Fig. 3b), this community was largely maintained through the colon until the stool, where the Firmicutes were reduced (Fig. 3b).

**Unique molecules from the microbiome.** Molecular networking paired with statistical analysis enabled identification of molecules unique or enriched between the two groups of mice. These included bile acids, flavonoids, triterpenoid saponins, and urobilins (Fig. S3-S5). The soyasaponins and flavonoids were prevalent, diverse and differentially abundant between the two groups of mice. These compounds were sourced from the mouse chow that had a dominant soybean component. A cluster of 76 connected nodes in the molecular network representing soyasaponins was found in both GF and SPF mice and their food pellets, but these clusters were enriched in nodes from the GI tract of GF mice (Fig. 3a). This molecular family contained a variety of unique soyasaponins all comprised of the core soyaspongenol triterpenoid backbone, but with different glycosylations and hydroxylations. Soysaponins were present throughout the GI tract of GF mice, including the stool sample, but in SPF mice they disappeared upon passage into the cecum (Fig. 3a). Conversely, there was a separate cluster only found in SPF mice that was annotated as soyasapogenols, which represent the triterpenoid backbone of soyasaponin without glycosylation (Fig. 3b). 3D-molecular cartography^2^ showed that soyasaponin I was abundant throughout the GI tract of GF mice, particularly the cecum, colon and stool, but was absent from these organs in SPF animals. In direct contrast, soyasapogenol was not found at all in GF animals, but was detected in the cecum of the SPF mice through to the stool. This differing presence of the glycone and aglycone forms indicates that cecal microbial activity was responsible for the metabolism of soyasaponin into soyasapogenol by removal of the saccharides (Fig. 3c-f). The abundance of soyasapogenol E (*m/z* 457.36) was then regressed against the microbiome data for significant associations between this metabolite and microbial operational taxonomic units (OTUs) (Bonferonni corrected p-value for 195 OTUs p < 2.6 x10^-4^). The Firmicute *Allobaculum* sp. (Pearson’s *r* = 0.491) was significantly correlated to the abundance of soyasapogenol E; the only cultured representative of this genus contains the β-glucosidase enzyme known to perform deglycosylation reactions of plant natural products^3^ (Table S1).

Microbiome breakdown of plant flavonoids was also observed (Fig. S4). In the mouse chow, glucuronides and aglycone flavones and isoflavones were detected, but not their sulfated forms. Because many isomeric forms of flavonoids exist that cannot be differentiated with our MS/MS methods, we focused on molecular changes in the predominant soybean isoflavonoids daidzein, genistein and glycitein, because they have characteristic MS/MS signatures^4^. In the GF mice, 3D-molecular cartography showed that the glucuronidated and sulfated isoflavonoids were detected throughout the GI tract from the stomach through to the stool, indicating they pass through the GI tract intact. In SPF mice, however, these same glucuronides and sulfides were undetectable in the distal GI tract. The aglycones were present in both the GF and SPF mice, but more abundant in the distal GI tract of GF animals (Fig. S4, Mann-Whitney U-test, p<0.05). Because the aglycones were detected in both groups, host and microbial enzymes (or chemical processes) could have been responsible for deglycosylation; however, the complete removal of the sugars and sulfates in the SPF mice indicated that the microbiota significantly enhanced this process. Furthermore, in the cecum of the SPF mice, the aglycone isoflavonoids were depleted and in some cases no longer detectable through to the stool samples, indicating that further metabolism of these compounds was occurring in the cecum and colon due to the presence of bacteria.

Detailed metabolism of plant flavonoids and triterpenoids was evident throughout the molecular networking data, particularly in the cecum and colon. The ability to not only visualize the location of this metabolism, but also identify specific chemical modifications of these compounds enabled a detailed understanding of microbiome metabolism of food molecules and other exogenous compounds in this study. Soyasaponins, triterpenoid glycosides found in soybeans and other legumes, were highly modified by the microbiota. These molecules have been associated with a variety of health and physiological effects including lowering cholesterol ^5^, anti-carcinogenic^6,7^, hepatoprotective^8^ and even antiviral activities ^9^. Mono- di- tri- and tetra-glycoside soyasaponins were all deglycosylated by the microbiome in the cecum, producing the common aglycone soyaspongenol despite the diversity and complexity of the parent glycone. It has been previously shown that the anti-carcinogenic properties of soyasaponins on human colon carcinoma cells only exist when these compounds are deglycosylated into soyasapogenols^7^. Thus, the beneficial effects of these compounds are likely dependent on the microbiota as the host cannot generate the aglycone with its own enzymes. Flavonoids were also altered by the murine microbiome through deglycosylation. The parent glycones and sulfated forms were absent from colonized animals, indicating complete microbiome-dependent removal of these accessory groups to produce the base flavonoid in the upper GI tract. In the lower GI tract, the aglycone isoflavonoids were absent or reduced in abundance, indicating either further metabolism of these plant natural products to other compounds by the microbiome or their microbiome-dependent absorption into the bloodstream^10,11^. Thus, the gut microbial community has a direct effect on the bioavailability of plant aglycone natural products and any benefits provided by them, which are believed to be extensive^12^. This location-specific metabolism of plant natural products seen in this study indicates that there may be potential to optimize the beneficial effects of these molecules by co-administration or enrichment with a probiotic in the region of the gut that best provides uptake of the aglycone.

**Bile acid metabolism.** Taurocholic acid (*m/z* 516.299), the major conjugated bile acid in mice^13^, was detected in all GI tract sections of GF mice as well as the liver and gallbladder. However, taurocholic acid was not detected beyond the cecum in colonized animals (Fig. S5). Instead, muricholic acid (*m/z* 409.295), the deconjugated form in mice, was present throughout the GI tract of SPF animals, including the stomach, liver and gallbladder. In GF mice, this metabolite was only detected in the liver, likely reflecting its role as the steroidal precursor of the major murine conjugated bile acids. Microbial specific secondary bile acids including deoxymuricholic acid (*m/z* 393.284) and deoxyketomuricholic acid (*m/z* 391.284) (Fig. S5) were found throughout the GI tract of SPF mice including the stomach, but were completely absent from GF mice. The abundant deoxymuricholic acid was regressed against the abundance of OTUs in the microbiome data in the samples where it was detected. A number of bacterial OTUs were significantly correlated with this molecule (Bonferonni corrected p-value p< 4.7 x 10^-4^). Unclassified members of the families Clostridiaceae (*r* = 0.746), Coprobacillaceae (*r* = 0.600), and Lachnospiraceae (*r* = 0.567) were significantly correlated with deoxymuricholic acid as well as a *Dehalobacterium* sp. (*r* = 0.643), *Coprococcus* sp. (*r* = 0.559), *Anaerotruncus* sp. (*r* = 0.559) and a *Blautia* sp. (*r* = 0.541).

Unique mass additions detected only in GF mice corresponded to acetylations (+42.045 Da) on taurocholic acids and tauroursocholic acids and a sulfation (+79.995 Da) of glycocholic acid. Their absence in colonized animals indicated that these chemical groups may have been removed by the microbiome. In the taurocholate-glycocholate molecular network, there was a cluster of five nodes unique to SPF mice without annotations in the GNPS libraries. The structures of the other two nodes unique to the microbiome in the glycocholic acid cluster remain unknown but show an addition of oxygen and an uncharacterized 50.968 Da to glycocholic acid (Fig. 4a).

**References**

1.	Tripathi, A. *et al.* Intermittent Hypoxia and Hypercapnia, a Hallmark of Obstructive Sleep Apnea, Alters the Gut Microbiome and Metabolome. *mSystems* **3,** e00020-18 (2018).

2.	Protsyuk, I. *et al.* 3D molecular cartography using LC–MS facilitated by Optimus and ’ili software. *Nat. Protoc.* **13,** 134–154 (2017).

3.	Michlmayr, H. & Kneifel, W. β-Glucosidase activities of lactic acid bacteria: mechanisms, impact on fermented food and human health. *FEMS Microbiol. Lett.* **352,** 1–10 (2014).

4.	Kuhn, F., Oehme, M., Romero, F., Abou-Mansour, E. & Tabacchi, R. Differentiation of isomeric flavone/isoflavone aglycones by MS2 ion trap mass spectrometry and a double neutral loss of CO. *Rapid Commun. Mass Spectrom.* **17,** 1941–1949 (2003).

5.	Lin, Y., Meijer, G. W., Vermeer, M. A. & Trautwein, E. A. Soy protein enhances the cholesterol-lowering effect of plant sterol esters in cholesterol-fed hamsters. *J. Nutr.* **134,** 143–8 (2004).

6.	Berhow, M. A., Wagner, E. D., Vaughn, S. F. & Plewa, M. J. Characterization and antimutagenic activity of soybean saponins. *Mutat. Res.* **448,** 11–22 (2000).

7.	Gurfinkel, D. M. & Rao, A. V. Soyasaponins: The Relationship Between Chemical Structure and Colon Anticarcinogenic Activity. *Nutr. Cancer* **47,** 24–33 (2003).

8.	Kinjo, J., Imagire, M., Udayama, M., Arao, T. & Nohara, T. Structure-Hepatoprotective Relationships Study of Soyasaponins I-IV Having Soyasapogenol B as Aglycone ^#^. *Planta Med.* **64,** 233–236 (1998).

9.	Kinjo, J. *et al.* Anti-herpes virus activity of fabaceous triterpenoidal saponins’). *Biol. Pharm. Bull.* **23,** 887–9 (2000).

10.	de Ferrars, R. M. *et al.* The pharmacokinetics of anthocyanins and their metabolites in humans. *Br. J. Pharmacol.* **171,** 3268–3282 (2014).

11.	van Duynhoven, J. *et al.* Metabolic fate of polyphenols in the human superorganism. *Proc. Natl. Acad. Sci. U. S. A.* **108 Suppl 1,** 4531–8 (2011).

12.	Cassidy, A. & Minihane, A.-M. The role of metabolism (and the microbiome) in defining the clinical efficacy of dietary flavonoids. *Am. J. Clin. Nutr.* **105,** 10–22 (2017).

13.	Swann, J. R. *et al.* Systemic gut microbial modulation of bile acid metabolism in host tissue compartments. *Proc. Natl. Acad. Sci. U. S. A.* **108 Suppl 1,** 4523–30 (2011).
