## Supplementary material for "Chemical Impacts of the Microbiome Across Scales Reveal Novel Conjugated Bile Acids": Methods

**Animals.** Germ-free (GF) C57Bl/6 mice were generated via caesarian section and microbiologically-sterile animals were cross-fostered by GF Swiss-Webster dams at the California Institute of Technology. GF animals were housed in open-top caging within flexible film isolators (Class Biologically Clean; Madison, WI) and maintained microbiologically sterile, confirmed via 16S rRNA PCR from fecal-derived DNA and culture of fecal pellets on Brucella blood agar or tryptic soy blood agar (Teknova; Hollister CA) under anaerobic and aerobic conditions, respectively. Conventionally-colonized specific pathogen free (SPF) mice were housed in autoclaved, ventilated, microisolator caging. All animals received autoclaved food (LabDiet Laboratory Autoclavable Diet 5010; St Louis, MO) and water *ad libitum*, were maintained on the same 12-hour light-dark cycle, and housed in the same room of the facility*.* All animal husbandry and experiments were approved by the California Institute of Technology’s Institutional Animal Care and Use Committee (IACUC). All animal dissections and sample collection for the GF and SPF mouse aspect of the study were carried out at University of California at San Diego under IACUC approval, protocol S00227M. For MRI imaging, a female, C57Bl/6 mouse, 8 weeks of age, was obtained from Jackson Laboratory and housed with food and water ad libitum. For metabolome and microbiome studies, four germ-free (GF) and four specific-pathogen-free (SPF) female 8-week-old C57Bl/6 mice were acquired from the California Institute of Technology’s vivarium. Samples of the food the animals were provided were also collected and analyzed (GF were fed LabDiet 5010 and SPF were fed LabDiet 5053, LabDiet, St. Louis, MO). An additional 24 male ApoE knockout mice in the C57BL/6J background raised for use in a study of hypoxia on the murine microbiome according to the methods of Tripathi et al. 2018^1^ were also analyzed in this study for the effects of high-fat-diet on bile acids. The fecal samples collected and data presented here were not published in that study.

**Human Sample Collection:** The unpublished work on samples of CF patients were collected from Rady’s Children’s Hospital in San Diego, CA using dual fecal swabs according to the procedure outlined in the American Gut Project^2^ under IRB approval #160034 and additional patients fecal samples and healthy controls were collected at Yale New Haven Hospital (New Haven, CT) under IRB approval 1206010476 according to the procedure outlined in^3^. Stool samples from patients with IBD were collected as part of the UCSD IBD Biobank under IRB #131487. Human infant fecal samples were collected at the University of Michigan under IRB #103575.

**3D Model Generation:**  A female, C57Bl/6 mouse, 8 weeks of age, was euthanized using carbon dioxide inhalation and then immediately brought to the UCSD Center for Functional MRI. The MRI images were acquired on a Bruker 7T/20 MRI scanner using a quadrature birdcage transceiver.  A 3D FLASH protocol with TE/TR=6 ms/15 ms and matrix size 128x64x156 was used, prescribing a field of view to match the body size. The dicom files from the mouse MRI were imported into the Invesalius software^4^. In Invesalius, the dicom files were visualized as stacked images through the axial, sagittal and coronal slices. Organs of interest were then traced in each slice according to their best visualization in the different viewpoints. The tracing was done using ‘create new mask’ feature in Invesalius using the manual edition mode. The brush feature was used to trace the outline of each organ of interest in the appropriate slice, stack by stack, until the entire organ was outlined through all slices in each orientation such that its outline was smoothed and did not bleed into other organs. Numerous iterations of this process led to the correct mapping of each organ through the MRI stacked images. The ‘Configure 3D surface’ feature was then used to translate the 2D stack tracings into a 3D image of each organ. This was completed for all organs sampled except for blood and skin samples, successively until an entire 3D-model of all organs was built. A fecal sample was also added through configuring a 3D-surface, but this was not a feature of the MRI. Blender (<https://www.blender.org/>) was used to smooth the model and color each organ differently, enabling better visualization of the different organs and organ systems (Fig. S1). Blood and skin samples were not mapped onto the model and a representative fecal sample was added after MRI modeling using Invesalius to allow mapping to a theoretical fecal sample.

**Sample Collection**.  Mice were euthanized via carbon dioxide asphyxiation. Prior to dissection, external sites including the skin (left and right flank), ears, mouth and feet were sampled using a cotton swab with vigorous contact for 5 seconds. Blood was collected via cardiac puncture using a 22-gauge needle and 1 ml syringe. Mice were then sterilely dissected under open flame using straight scissors and fine forceps that were cleaned with 70% ethanol (v/v) between handling of each organ. Sections of each organ were made using sterile razor blades, with the number of sections listed in Table S1.The following organs were dissected: Adrenal gland, bladder, brain, cecum, cervix, colon, duodenum, esophagus, foot, gall bladder, heart, ileum, jejunum, kidney, liver, lung, ovaries, spleen, stomach, thymus, trachea, uterus and vagina. Four stool samples were also collected from each group of mice, although it is not known which mouse produced which stool sample. Sample collection for the additional published murine studies were completed according to^1,5^. In addition, fecal samples were collected from mice fed a high-fat diet starting at 10 weeks and compared to animals fed the control normal chow diet according to the methods of^1^. Data from^1^ was previously published, but the samples used in this study were not published as part of that manuscript.

**Sample Processing:** All samples were contained in 2 ml sterile Eppendorf® Biopur® Safe-Lock tubes, wet tissue mass recorded, and then flash frozen. For the swab samples, the wooden end of the swab was cut off with scissors and 1 ml of PBS was added. All of the non-swab samples were then diluted in a 1/10 mass/volume in sterile phosphate buffered saline. A Qiagen (Qiagen Inc., Valencia, CA) 5 mm stainless steel bead was added to each tube and the samples were homogenized in a Qiagen TissueLyzer II homogenizer at a frequency of 20/s for 5 min. After homogenization two aliquots of 50 μl of the homogenate or PBS/swab mix was added to separate 96-well deep well plates, one for metabolite extraction and one for DNA extraction. Metabolites were extracted from the samples in the 96-well deep well plate by adding 200 μl of LC-MS grade 70% methanol in LC-MS grade water. Samples were left to extract overnight at 4°C and then spun down to pellet debris in a 96-well plate Sorvall® Legend centrifuge at 2500 rpm for 1 minute. DNA was extracted using extraction according to protocols benchmarked for the Earth Microbiome Project (EMP) found here: <http://www.earthmicrobiome.org/emp-standard-protocols/>^6,7^.

**LC-MS/MS Mass Spectrometry:** A 50 μl aliquot of the extracted sample in methanol was added to a 96-well plate and diluted with 150 μl of LC-MS grade methanol containing 2 μl of ampicillin MS internal standard. The chromatographic separation was conducted on a ThermoScientific UltraMate 3000 Dionex UPLC system (Fisher Scientific, Waltham, MA USA) with eluent subsequently electrospray ionized and analyzed with a Bruker Daltonics^®^ MaXis qTOF mass spectrometer (Bruker, Billerica, MA USA). Metabolites were separated using a Kinetex 2.6 μm C18 (30 x 2.10 mm) UPLC column containing a guard column. Mobile phases A 98:2 and B 2:98 ratio of water and acetonitrile, respectively, containing 0.1% formic acid and a linear gradient from 0 to 100% for a total run time of 840 s at a flow rate of 0.5 mL min^-1^ were used. The mass spectrometer was calibrated daily using Tuning Mix ES-TOF (Agilent Technologies) at a 3 mL min^-1^ flow rate. For accurate mass measurements, lock mass internal calibration used a wick saturated with hexakis (1H,1H,3H- tetrafluoropropoxy) phosphazene ions (Synquest Laboratories, *m/z* 922.0098) located within the source. Full scan MS spectra (*m/z* 50 – 2000) were acquired in the qTOF and the top ten most intense ions in a particular scan were fragmented using collision induced dissociation at 35 eV for +1 ions and 25 eV for +2 ions in the collision cell. A data dependent automatic exclusion protocol was used such that an ion was fragmented upon its first detection, then fragmented twice more, but not again unless its intensity was 2.5x the previous fragmentation. This exclusion method was cyclical, being restarted after every 30 seconds. Mass spectrometry data for the mice fed a high-fat diet compared to normal chow for 10 weeks was generated separately from this study on a ThermoScientific® qExactive® mass spectrometer according to the procedure of ^1^.

**Metabolomics Data Processing** **and Analysis.** Each LC-MS/MS file in the Bruker format (.d) was converted to .mzXML format using the Bruker® DataAnalysis ‘Process with Method’ batch script. Lock mass calibration was applied during conversion to aid in mass accuracy. The .mzXML files were uploaded to the UCSD MassIVE data storage server for GNPS analysis. The entire dataset is publically available and found under the ID MSV000079949. In addition, the area under curve feature abundances were calculated in batch for all files using the Optimus software based on the OpenMS feature finding algorithms^8^. The Optimus parameters were as follows: m/z tolerance 15.0 ppm, noise threshold of 3000, retention time tolerance of 20 s, intensity factor compared to blanks at 3.0, and a feature observation rate of 0.01. The data was then trimmed to contain information only from 60 s to 550 s of the run during the linear gradient. The feature abundances were normalized to the total ion abundance in each sample for all statistical analysis. For organ-by-organ analysis the features present in individual organs were extracted as separate buckettables and any features not present at all in a particular organ were removed. Data generated with the ThermoScientific qExactive mass spectrometer from the HFD study^1^, was processed using the mzMine software^9^ and the feature table was normalized to the total ion current. Parameters were as follows: MS^1^ minimum threshold of 10000 counts, MS^2^ threshold of 5000 counts, a mass tolerance of 0.03Da and retention time tolerance of 0.2 min. The data was deconvoluted, deisotoped and filtered for compounds present in at least 3 samples. This additional metabolomics dataset is publically available under MassIVE ID MSV000082480.

       	Molecular networking was performed on GNPS with the GF and SPF mice separated from each other and from blank or quality control samples using the group-mapping feature. The molecular networking and MS-cluster parameters were as follows: parent and fragment ion mass tolerance 0.05 Da, minimum cosine score of 0.7, minimum matched fragment ions of 4, and a minimum cluster size of 4 (to minimize detection of more rare nodes found in few samples). The library search parameters of the molecular networking search were a minimum-matched peaks of 4 and a cosine score of 0.65. Any library hits from the results were inspected directly between the spectrum and query and are considered level two according to the metabolomics standards consortium guidelines^10^. The estimated false discovery rate (FDR) for spectral matching is 4.1% under our search parameters^11^. A link to the full data molecular network used for statistical analysis is available here <https://gnps.ucsd.edu/ProteoSAFe/status.jsp?task=9ea760fb819449d7bc7aca8fec07bd8d>.

Analysis of the mass shifts and chemical transformations between nodes was done using the method of^12^. Briefly, all nodes unique to either GF or SPF were searched for an edge connection to a node from one or the other groups (GF to SPF, SPF to GF, GF to shared or SPF to shared). In each instance, the mass gain or loss, relative to the unique node, was recorded along with the spectral count for each node. Mass differences were binned into known molecular modifications within a 0.003 Da window as described from MeMSChem with the addition of unique modifications in this dataset, such as saccharides. The spectral counts for each modification were counted and plotted as the relative abundance to all modifications and as total spectral counts for each modification.

**16S rRNA Gene Sequencing:** On all murine samples collected both GF and SPF and control samples of solutions and swabs underwent DNA extraction, 16S rRNA gene variable region 4 (V4) PCR and amplicon preparation for sequencing according to protocols benchmarked for the Earth Microbiome Project (EMP) found here: <http://www.earthmicrobiome.org/emp-standard-protocols/> ^7,13^. The microbiome data was processed through the Qiita software (qiita.ucsd.edu) The data was demultiplexed and then rarified at a sequence sampling depth of 500 before processing using the closed reference OTU picking method. The resultant .biom files were used for downstream analysis. The microbiome data is available at (<https://qiita.ucsd.edu/> study ID: 10801/).

**3D Mapping in *‘ili*:** Metabolomics and microbiome data were mapped onto the 3-D mouse model by recording the location of the sampling and orientation of each sample in the model according to the methods described in^14^. Some organs only contained one sample (bladder, blood, cervix, gall bladder and thymus) all other organs contained 2-6 samples and the actual location of the dissected sample was mapped to the appropriate point representing that same sample in the 3D model. The point mapping was done using the GeoMagic® Wrap software. The full .stl model of the laboratory mouse was loaded into GeoMagic Wrap and the location of each sampling point was selected with the ‘points’ tool. The x,y,z coordinate information in the model from all points was then exported as a .csv file for matching to its representative sample in the metabolomics or microbiome data. Sub models of different organ systems were also created in the same manner to aid visualization, such as the GI tract and liver. Mapping to these models was done as described for the full model. For ‘*ili* visualization the matching samples for the 4 GF and 4 SPF mice were averaged and a new feature or OTU table created based on these mean abundances. This feature table was then matched to the x,y,z coordinates from the model according to the correct sample. This OTU or metabolite feature table was then uploaded into the *‘ili* software simultaneously with the mouse model. This enabled automatic mapping of the abundance of a microbial or metabolite variable to the point representing its collection location in the GF and SPF mouse 3D-model. Visualization in ‘*ili* was done using a linear scale with the ‘viridis’ color map and automatic min/max mapping was selected.

**Statistical Analysis:** The microbiome .biom table and metabolome feature table were analyzed using principal coordinate analysis after calculation of a distance matrix between all samples. The microbiome distance matrix was generated using the weighted UniFrac distance^15^ in Qiita (qiita.ucsd.edu) and the metabolomic distance was calculated using the Bray-Curtis dissimilarity. The resulting distance matrix was visualized using PCoA and each sample highlighted by either GF/SPF or organ source. Beta-diversity of the microbiome data was calculated on non-rarified data to enable visualization of GF and sterile samples which had a low numbers of 16S rDNA gene reads. All other statistical analysis was completed on the rarified data.

       	To determine the number of unique metabolites between GF and SPF in each organ molecular networks were built with the same above parameters for samples from each of the 29 organs. The molecular networking data was then downloaded from GNPS and the source of each node as GF or SPF was tabulated. A spectrum was considered unique to either class of mice only if it was detected in at least 3 out of 4 individual mice sampled per category. Each instance of these unique nodes was counted and reported as a percentage of the total number of nodes from each organ and as the total number of nodes per organ to visualize abundance.

       	To visualize the effect of the GF or SPF classification on the gut metabolomic data a random forests classification was run on all GI tract samples (including the esophagus) and the variable importance for classification of each metabolite was determined. The 30 most differentially abundant metabolites according to their variable importance were then visualized using a stacked bar graph where the 30 metabolites summed to 100%. This enabled visualization of the changes in the most differential metabolites through the GI tract. In addition, the most abundant 24 metabolites (normalized abundance) were also presented in a stacked bar graph to visualize the change in these molecules through the GI tract. The mean Shannon-Weiner index of diversity was calculated on the entire metabolome from each GI tract associated sample using the R statistical software. The mean Shannon-Weiner diversity was then shown the two groups of mice through the GI tract. The Mann-Whitney U-test was used to determine a statistically significant difference (p< 0.05) between the Shannon diversity of each GI tract sample collected at the same location between the GF and SPF mice. The microbiome diversity was calculated using the Faith’s phylogenetic diversity index in the Qiita software and mean diversity between the four individual mice was presented only for the SPF mice.

Tests of the differential abundance of the novel bile acids between mice fed antibiotics or high fat were done using the Mann-Whitney U-test with a significance level of p<0.05.

**Novel Bile Conjugates Validation Experiments.** To validate the synthetic standards of the tyrosine, phenylalanine, leucine and isoleucine cholic and muricholic acids, the compounds were dissolved in methanol, diluted to approximately 5 μM and run on the LC-MS/MS method described above. The data is publically available under MassIVE ID: MSV000082467. Retention times and MS/MS spectra were analyzed to verify the molecular characteristics. To determine the approximate concentration of phenylalanocholic acid in the murine GI tract a ileal sample from a GF mice was spiked with standard curve of concentrations of pure phenylalanocholic acid (non-murine form). Final concentrations of 100 μM, 25 μM, 5 μM, 1 μM, 0.1 μM and 0.02 μM, were directly added to the extracted ileal sample and analyzed with mass spectrometry using the same methods as described above. A standard curve of these concentrations was calculated by plotting the known concentrations to their corresponding area-under-curve (AUC) abundance of the phenylalanocholic acid peak. The same AUC abundance was then captured for each sample positive for the molecule in the colonized mice. The concentration in the murine samples was then calculated based on the concentrations of the standard curve. Because isoleucine and leucine cannot be distinguished with MS/MS data, we analyzed the synthetic isoleucocholic acid standard and leucocholic acid standard on an extended gradient HPLC column. The two standards were injected with the jejunum3 sample from mouse SPF2 and subjected to a 40% LC gradient of the same solvents described above with ramp to 40% solvent B at 3 minutes followed by 22 min of ramping to 100% B and then wash steps. The MS/MS method was identical to that described above and retention time differences were recorded between the two chemical standards and the murine sample. To determine whether the base bile acid was either cholic or muricholic acids, the muricholic forms were synthesized according to the supplementary methods in place of cholic acids and all 3 amino acid conjugates of each bile acid backbone were analyzed using the original LC-MS/MS with sample SPF2 jejunum 3, which contained the same molecules detected in the murine gut. Retention time analysis was used to identify whether each molecule in the mouse sample was either muricholic or cholic acid forms.

**Mining Public Data on GNPS**. The single spectrum search feature in GNPS (<https://bit.ly/2M9KyyD>) that allows one to search all public data akin to BLAST allows one to search public sequence information. This is a new feature in GNPS and is used here for the first time but is accessible to anyone (https://bit.ly/2M9KyyD) and molecular explorer from GNPS (Wang et al. 2016) were used to search for the unique amino acid conjugated bile acids in publicly available data. In datasets with a positive hit, the source organism and % of samples positive for each compound was recorded. Two datasets comprised of LC-MS/MS data analyzed on a Bruker Maxis qTOF from fecal swabs of CF patients (massive IDs MSV000079134 and MSV000082406) were further analyzed according to the metadata of the studies as pancreatic sufficient, insufficient or samples from healthy individuals. The presence of an MS/MS spectrum for each of these classes was tabulated by individual and reported as the percent of subjects positive for each molecule in each class.

**References**

1.	Tripathi, A. *et al.* Intermittent Hypoxia and Hypercapnia, a Hallmark of Obstructive Sleep Apnea, Alters the Gut Microbiome and Metabolome. *mSystems* **3,** e00020-18 (2018).

2.	McDonald, D. *et al.* American Gut: an Open Platform for Citizen Science Microbiome Research. *mSystems* **3,** e00031-18 (2018).

3.	Cullen, T. W. *et al.* Antimicrobial peptide resistance mediates resilience of prominent gut commensals during inflammation. *Science (80-. ).* **347,** 170–175 (2015).

4.	Amorim, P., Moraes, T., Silva, J. & Pedrini, H. in 45–54 (Springer, Cham, 2015). doi:10.1007/978-3-319-27857-5_5

5.	Shalapour, S. *et al.* Inflammation-induced IgA+ cells dismantle anti-liver cancer immunity. *Nature* **551,** 340–345 (2017).

6.	Caporaso, J. G. *et al.* QIIME allows analysis of high-throughput community sequencing data. *Nat. Methods* **7,** 335–6 (2010).

7.	Caporaso, J. G. *et al.* Ultra-high-throughput microbial community analysis on the Illumina HiSeq and MiSeq platforms. *ISME J.* **6,** 1621–4 (2012).

8.	Kenar, E. *et al.* Automated Label-free Quantification of Metabolites from Liquid Chromatography–Mass Spectrometry Data. *Mol. Cell. Proteomics* **13,** 348–359 (2014).

9.	Pluskal, T., Castillo, S., Villar-Briones, A. & Orešič, M. MZmine 2: Modular framework for processing, visualizing, and analyzing mass spectrometry-based molecular profile data. *BMC Bioinformatics* **11,** 395 (2010).

10.	Sumner, L. W. *et al.* Proposed minimum reporting standards for chemical analysis Chemical Analysis Working Group (CAWG) Metabolomics Standards Initiative (MSI). *Metabolomics* **3,** 211–221 (2007).

11.	Scheubert, K. *et al.* Significance estimation for large scale metabolomics annotations by spectral matching. *Nat. Commun.* **8,** 1494 (2017).

12.	Hartmann, A. C. *et al.* Meta-mass shift chemical profiling of metabolomes from coral reefs. *Proc. Natl. Acad. Sci. U. S. A.* **114,** (2017).

13.	Caporaso, J. G. *et al.* Global patterns of 16S rRNA diversity at a depth of millions of sequences per sample. *Proc. Natl. Acad. Sci. U. S. A.* **108 Suppl,** 4516–22 (2011).

14.	Protsyuk, I. *et al.* 3D molecular cartography using LC–MS facilitated by Optimus and ’ili software. *Nat. Protoc.* **13,** 134–154 (2017).

15.	Lozupone, C. & Knight, R. UniFrac: a new phylogenetic method for comparing microbial communities. *Appl. Environ. Microbiol.* **71,** 8228–35 (2005).
